# The landscape of T cell epitope immunogenicity in sequence space

**DOI:** 10.1101/155317

**Authors:** Masato Ogishi, Hiroshi Yotsuyanagi

## Abstract

The existence of population-wide T cell immunity is widely recognized for multiple pathogen-derived immunodominant epitopes, despite the vast diversity and individualized nature of T cell receptor (TCR) repertoire. We thus hypothesized that population-wide epitope immunogenicity could be probabilistically defined by exploiting public TCR features. To gain a proof-of-concept, here we describe a machine learning framework yielding probabilistic estimates of immunogenicity, termed “immunogenicity scores”, by utilizing features designed to mimic thermodynamic interactions between peptides bound to major histocompatibility complex (MHC) and TCR repertoire. Immunogenicity score dynamics among observed and computationally simulated single amino acid mutants delineated the landscape of position- and residue-specific mutational impacts, and even quantitatively estimated escaping potentials of known epitopes with remarkable positional specificity. This study illustrates that the population-wide aspect of adaptive immunity is predictable via non-individualized approach, possibly indicating antigen-guided convergence of human T cell reactivity.

## Introduction

T cell epitopes bound to major histocompatibility complex [MHC; also called the human leukocyte antigen (HLA) in humans] molecules activate T cells to initiate subsequent immunological orchestration (Garcia et al., 1999; Grakoui et al., 1999; Rudolph et al., 2006). MHC class I (MHC-I) molecules typically present 8- to 11-aa peptides generated through proteasomal cleavage of intracellular proteins to activate CD8+ cytotoxic T lymphocytes (CTLs), whereas MHC class II (MHC-II) molecules with an open-ended binding groove accommodate peptides of more variable length derived from endocytosed proteins to activate CD4+ T helper (Th) cells (Neefjes et al., 2011). Evidence suggests that not all peptides presented on MHC molecules are immunogenic, *i.e.*, trigger functional T cell activation (Bjerregaard et al., 2017; Newell et al., 2013; Sette et al., 1994). T cells recognize peptide-MHC (pMHC) complexes by their TCRs, most predominantly via complementarity determining region 3 (CDR3) loops (Borg et al., 2005; Tian et al., 2007; Zhong et al., 2013). TCR CDR3 sequences are exceedingly diversified (known as TCR repertoire) through gene segment rearrangement and in rare circumstances through somatic hypermutation (Cheynier et al., 1998). However, contrary to the general notion that TCR repertoire is highly stochastic and individualized, accumulating evidence suggests that the biased generation of TCR repertoire due to non-random rearrangement and positive/negative thymic selections is thought to result in an extensive utilization of a limited subset of possible sequences and larger-than-expected overlaps of repertoires between individuals knowns as public clonotypes (Britanova et al., 2014; Madi et al., 2014; Ndifon et al., 2012; Robins et al., 2010; Shugay et al., 2013). Moreover, even distinct TCR repertoires can recognize identical immunodominant epitopes (Chen et al., 2017). Finally, population-wide immunity to some pathogen-derived epitopes has been proven and even utilized clinically, exemplified by the interferon-gamma release assay for tuberculosis (Horvati et al., 2016; Udhayakumar et al., 1995). These observations led us to the hypothesis that population-wide epitope immunogenicity could be probabilistically defined, and public TCR usage pattern might serve as a probe. In line with this hypothesis, residue hydrophobicity at TCR contact residues has been associated with immunogenicity, although the predictive capability was not satisfactory (Chowell et al., 2015). In this context, here we propose that a computational framework mimicking the intermolecular interactions between pMHC complexes and public human TCR clonotypes, termed TCR-peptide contact potential profiling (CPP), generates probabilistic estimates of immunogenicity, termed immunogenicity scores, which effectively recapitulate essential characteristics of T cell immunity and enable quantitative assessment of potential mutational impacts on the dynamics of immunogenicity, *i.e.*, escaping, in a position-specific context.

## Result

### Concepts and analysis workflow

To gain a proof-of-concept, we first compiled epitope datasets comprising 21,162 8- to 11-aa HLA-I-restricted and 31,693 11- to 30-aa HLA-II-restricted peptide sequences from various sources (Figure 1, Figure S1 and Data S1) (Calis et al., 2013; Chowell et al., 2015; Fleri et al., 2017; Kuiken et al., 2005; Lata et al., 2009; Llano et al., 2013; Olsen et al., 2017; Reche et al., 2005; Tung et al., 2011). We defined positivity as the existence of at least one positive functional T cell assay result. A total of 6,957 and 16,642 HLA-I and HLA-II-restricted epitopes met the criteria. Next, we extracted 23,006,555 unique TCR β chain CDR3 (CDR3β) sequences from TCR repertoire datasets derived from healthy individuals using MiXCR software (Bolotin et al., 2015). We obtained 191,326 unique CDR3β sequences identified in at least 22 out of 206 different datasets to construct fragment libraries for the CPP feature computation.

**Figure 1.**
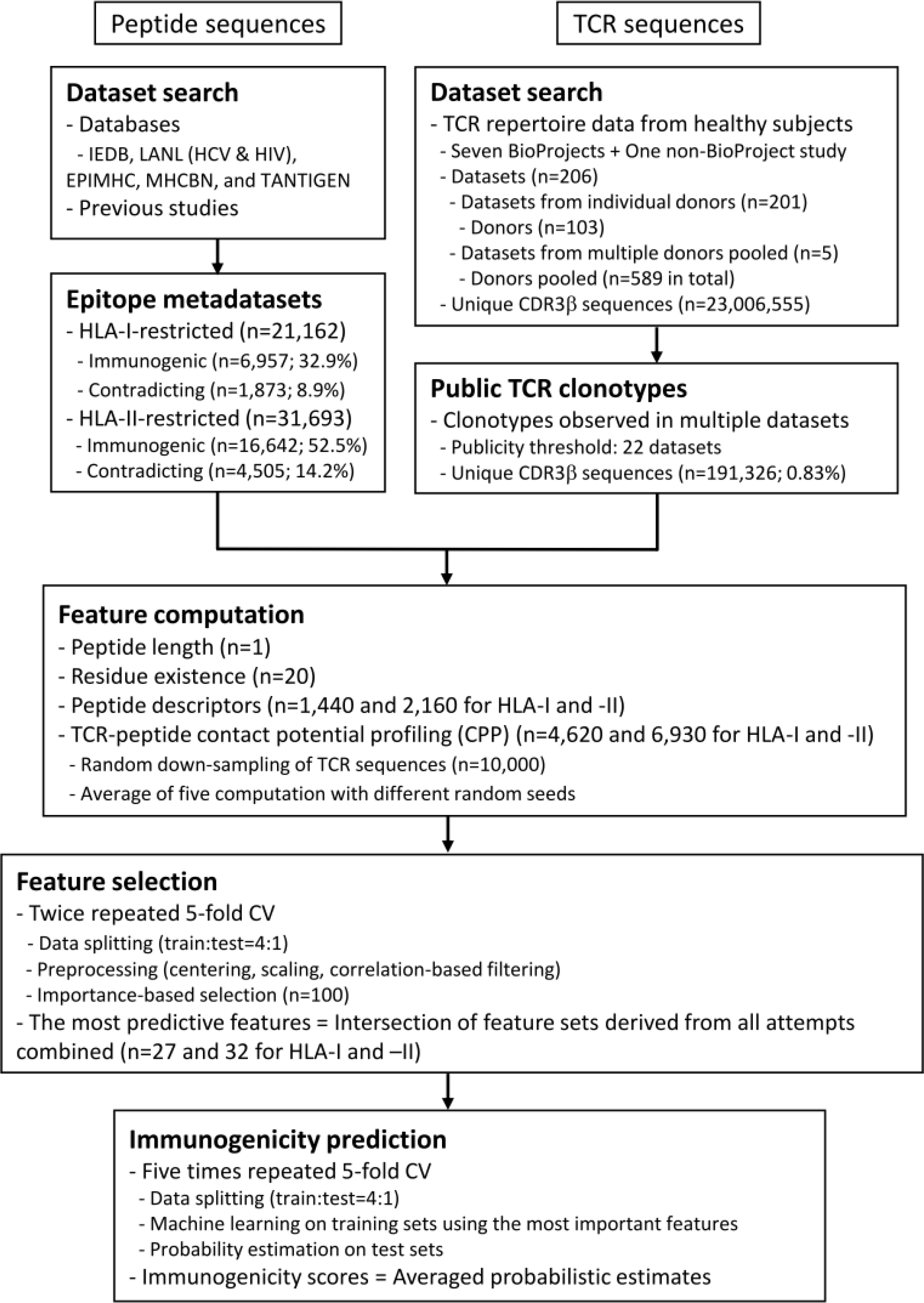
Data analysis workflow. The workflow for generation of the probabilistic estimates of epitope immunogenicity, termed immunogenicity scores, is shown. All datasets were retrieved from public sources. CV, cross-validation. See also Figures S1 and S2, Data S1, and Materials and Methods.

The concept of CPP relies on the hypothesis that public TCR repertoire is biased toward immunodominant epitopes through evolutional adaptation and thymic negative selection, and therefore could be used to probe peptides with the highest immunogenic potentials (Figure 2A) (Blackman et al., 1990; Madi et al., 2017; Miles et al., 2011; Robins et al., 2010). However, because of the sequence-level and structural diversity of TCR and pMHC, it is unfeasible to comprehensibly characterize every single TCR-pMHC interaction experimentally. To circumvent this problem, we instead designed the CPP framework with the following simplifications. First, we focused on CDR3β loop because this region has the highest genetic variability and is primarily responsible for the interactions with peptides presented onto the MHC grooves, whereas more conserved CDR1 and CDR2 loops typically interact with MHC a-helices (Bridgeman et al., 2012; Cole et al., 2014; Garcia et al., 1999; Rudolph et al., 2006). Second, we assumed that initial intermolecular interaction requires only a small portion of the CDR3β loop that could be approximated to linear structure, based on the low affinities (*K*_*D*_ = 0.1 ~ >500 μM) compared to other immunoglobulin-like molecules including antibodies, the relatively flat interfaces, and the substantial conformational changes induced upon recognition of pMHC complexes (Boniface et al., 1999; Garcia et al., 1998; Hennecke and Wiley, 2001; Lee et al., 2004). Indeed, analysis of known TCR-pMHC complex structures using PRODIGY (Vangone and Bonvin, 2015) suggested that around 50% and more than 80% of the intermolecular contacts are limited within the 3-aa and 5-aa ranges, respectively, in both peptides and CDR3β regions (Figure 2C and Data S2). Finally, we approximated the “best-match” problem between peptides and TCR repertoire to pairwise sequence alignment problem, where the alignment scores are interpreted to reflect the contact energy potentials (Figure 2D).

**Figure 2.**
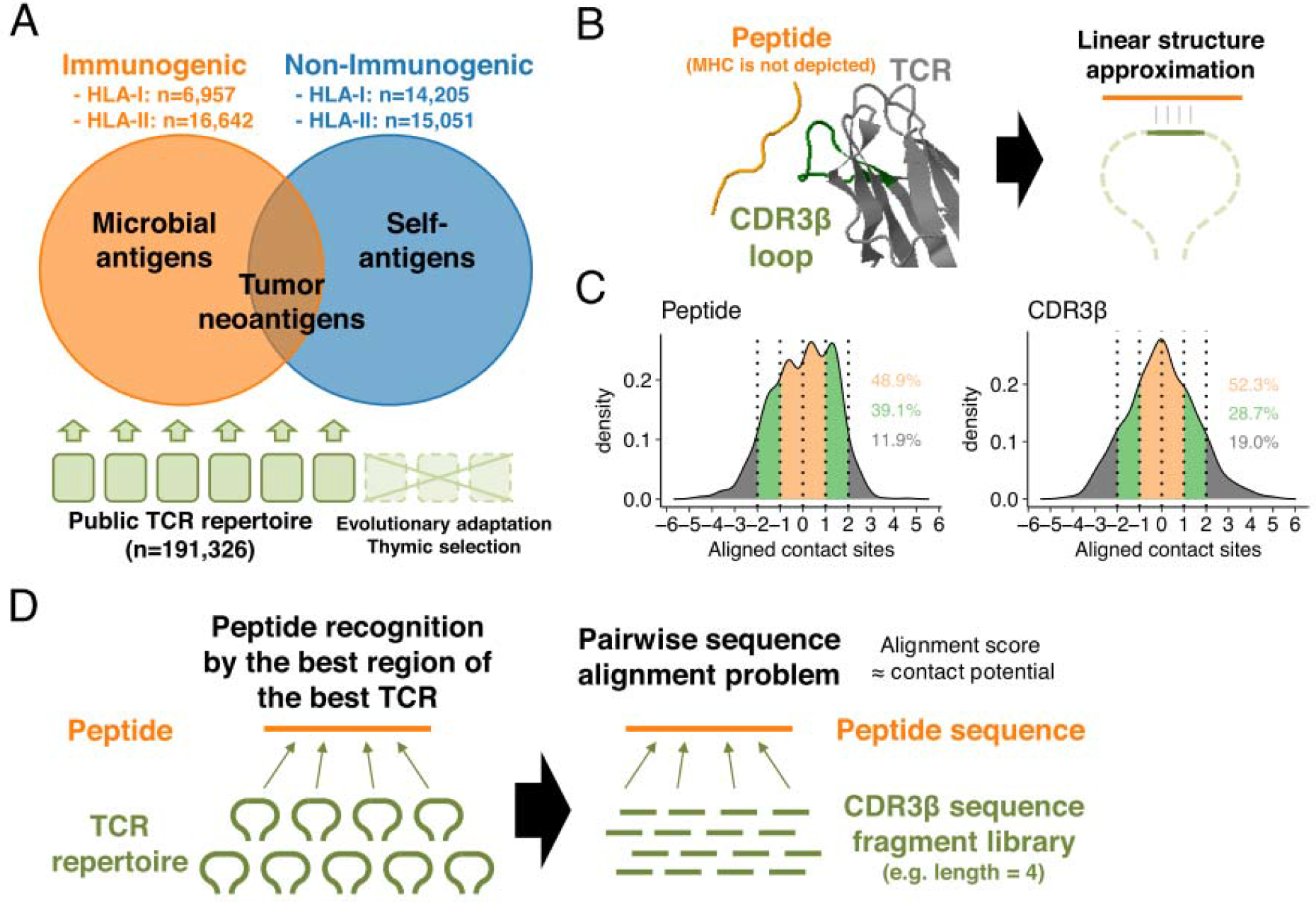
TCR-peptide contact potential profiling. (A) Schematic of antigen-guided TCR repertoire convergence hypothesis. We formulate the concept of TCR-peptide contact potential profiling (CPP) to identify population-wide immunogenicity by exploiting public TCR repertoire inherently biased toward common pathogen antigens but away from self-antigens due to genome-level evolution and postnatal thymic selection. (B) Schematic of the linear contact model. The underlying hypothesis is that only a small portion of the CDR3β loop structure recognizes an epitope. The structure of HIV reverse-transcriptase epitope (TAFTIPSI) and cognate TCR CDR3β chain is shown as an example (PDB ID: 4MJI). (C) Mean-centered contact density distributions determined from 51 unique peptide-WT TCR structures. Inset numbers indicate the proportions of contacts falling into the color-specified ranges. (D) Schematic of the approximation of best-matching region problem to pairwise sequence alignment problem. See also Data S2, and Materials and Methods.

### TCR contact features essential for epitope immunogenicity prediction

We generated CPP features using public clonotype sequences, as well as other features based on peptide physicochemical properties (Figure 1). Feature selection yielded the 32 and 27 most predictive features for MHC-I and MHC-II predictions, respectively (Figure S2 and Data S3). Only nine of 32 and none of 27 features were derived from peptide physicochemical descriptors, highlighting the indispensable contributions of CPP features on immunogenicity prediction. Common parameter usage patterns for CPP features were observed in MHC-I and MHC-II (Figure 3A-C). Notably, features derived from short fragments (*i.e.*, 3-aa and 4-aa) and the longest fragment (*i.e.*, 8-aa and 11-aa in MHC-I and MHC-II, respectively) had higher cumulative importance values (Figure 3A). Meanwhile, skewness- and kurtosis-derived features showed markedly higher importance, indicating that distinct repertoire-wide contact potential distributions are the hallmark of immunogenicity (Figure 3B). The inverted version of BETM990101 (termed “BETM990101inv”) (Betancourt and Thirumalai, 1999) and other six AAIndex scales of the highest importance for both MHC-I and MHC-II (Figure 3C). To test the hypothesis that selected AAIndex scales may reflect thermodynamic aspects of TCR-peptide interactions, we analyzed 82 TCR-peptide complexes with experimentally determined affinities collected from literature (Data S2) through a modified framework which we termed single-TCR contact potential profiling (sCPP). We found that BETM990101inv and other six AAIndex scales important for immunogenicity prediction also generated sCPP features correlating with affinities (Figure 3D and E and Table S1). Sequence permutations significantly undermined the observed correlations (Figure 3F). Collectively, these observations support the notion that our CPP framework recapitulates the essential thermodynamic properties of TCR recognition of pMHC complex which governs both MHC-I and MHC-II systems.

**Figure 3.**
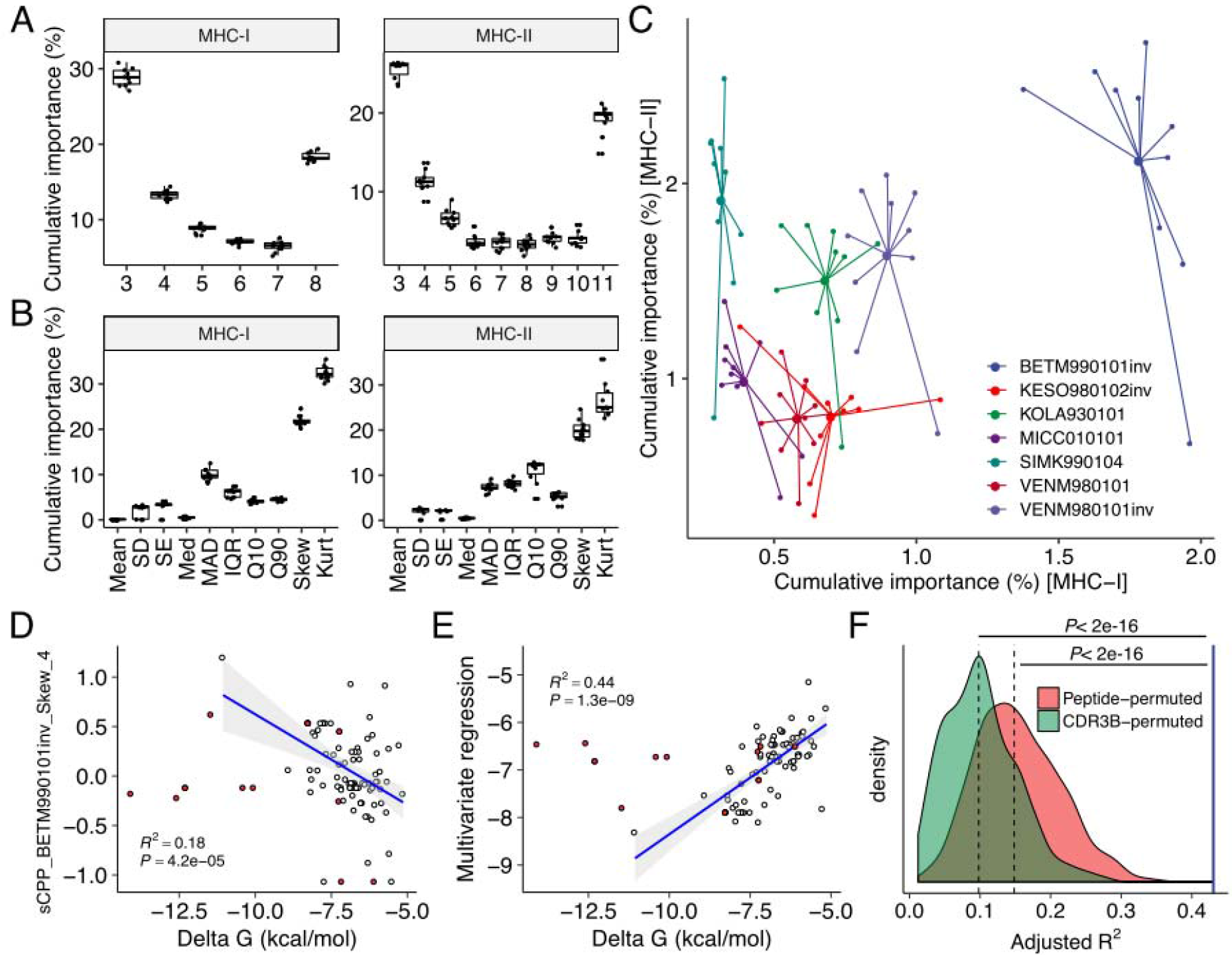
Contact potential profiling parameters linked epitope immunogenicity and TCR affinity. (A-C) Feature importance estimates from ten iterations. (A and B) Importance estimates for all CPP features aggregated by (A) fragment length and (B) descriptive statistics. (C) Importance estimates for the most predictive 21 and 26 CPP features for MHC-I and MHC-II, respectively, aggregated by their source AAIndex scales. The seven most predictive AAIndex scales selected both in MHC-I and MHC-II are shown. (D-F) Regression analysis against individual TCR affinities. Single-TCR contact potential profiling (sCPP) was conducted for 82 wildtype (WT) TCR-peptide complexes. Thirteen mutant (MT) TCRs were excluded from correlation analysis and only shown for visualization purposes (red points). Adjusted squaredPearson’s correlation coefficients are presented. (D) Representative univariate regression. (E) Multivariate regression with five sCPP features derived from immunogenicity-predicting AAIndex scales. (F) Permutation experiments with 1,000 iterations. Multivariate regression was performed as in (E) using sCPP features computed from either randomly permuted peptide sequences or CDR3β sequence fragments. The blue line represents the non-permuted regression result. Dashed lines represent medians. See also Figure S2, Data S2 and S3, and Table S1.

### Probabilistic estimation of epitope immunogenicity

We applied machine learning techniques to convert these most predictive features into a unidimensional scale of population-wide immunogenicity by averaging the probability estimates from five times repeated five-fold cross-validations (CVs). We termed this scale as “immunogenicity scores” (Data S4). The estimates were reasonably consistent across CVs; the normalized standard deviations distributed with medians and interquartile ranges (IQRs) of 8.3% (2.3% to 14.3%) and 5.0% (1.4% to 8.5%) in MHC-I and MHC-II, respectively. Addition of MHC binding prediction results during machine learning only marginally improved the predictive performance for MHC-I and had almost no effect for MHC-II. Both reduction of features utilized during machine learning and removing peptides with high homology from the dataset resulted in only a moderate decrease in predictive performance (Figure 4A and Figure S3). Contrastingly, the MHC-I immunogenicity prediction tool (Calis et al., 2013) showed only marginal predictive power on our dataset, with the area under the ROC curve (AUC) of 54% (Figure S4), and there remains no existing tool for MHC-II to compare. Immunogenicity scores stably differentiated epitopes from non-immunogenic binders across various subsets of the datasets, including data obtained from autologous *ex vivo* assays (Figure 4B). Extrapolation of human immunogenicity classifiers to non-human peptide datasets yielded estimates with slightly lower predictive performances (Figure 4C and Figure S5), although the dearth of data prevents us from drawing a solid conclusion on MHC-II-restricted primate epitopes.

**Figure 4.**
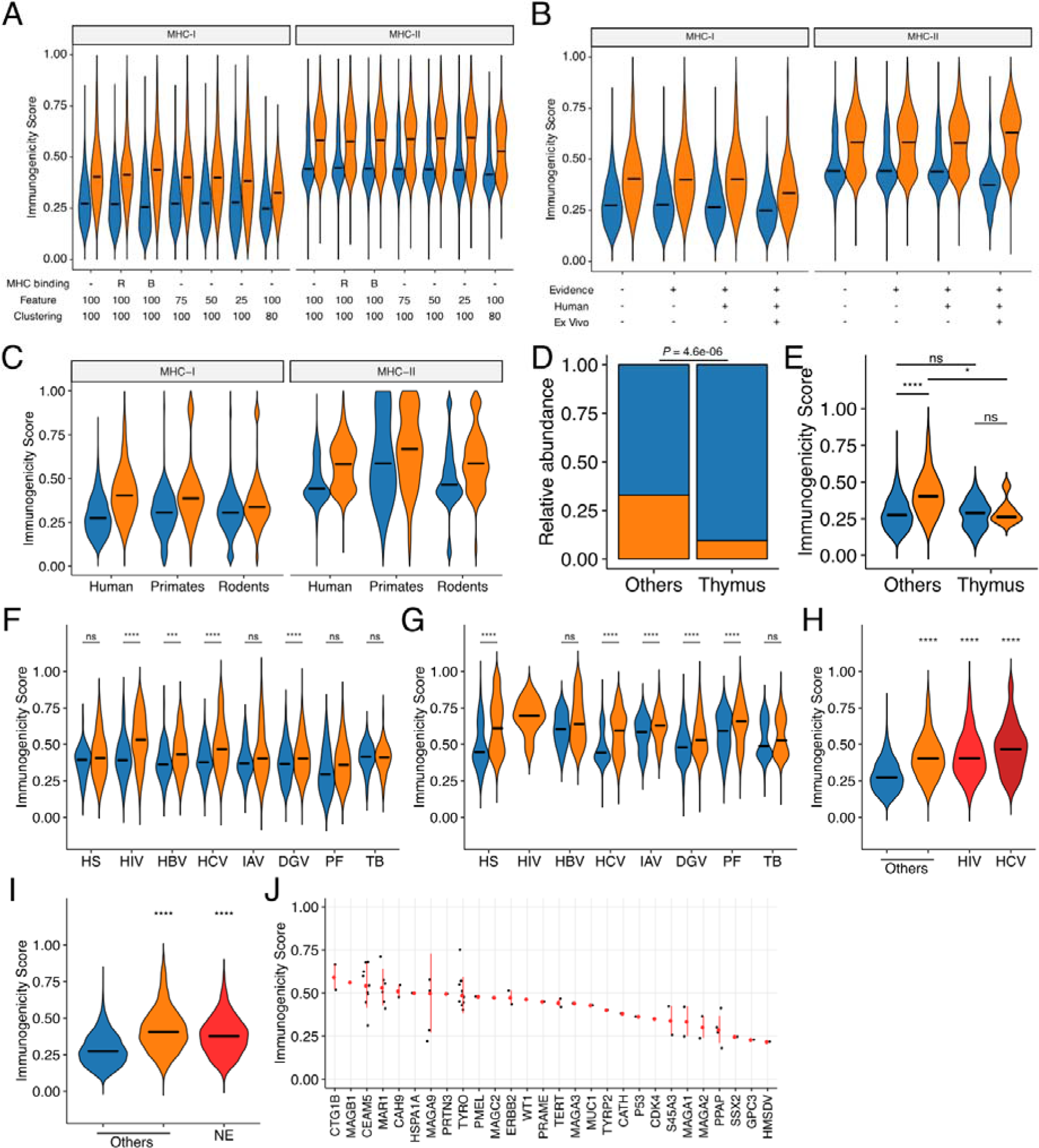
Immunogenicity prediction. Averaged probabilistic estimates from iterative machine learning were termed “immunogenicity scores.” (A) Immunogenicity scores generating under various conditions for machine learning processes. MHC binding: incorporation of MHC binding prediction results in machine learning. R, percentile ranks; B, binding strength categories. Feature: the cutoffs for feature selection. The numbers of features actually retained after feature selection under the cutoff of 100, 75, 50, and 25 were 32, 26, 19, and 7 for MHC-I, and 27, 23, 21, and 12 for MHC-II, respectively. Clustering: Elimination of highly similar peptide using the IEDB Epitope Cluster Analysis Tool (http://tools.iedb.org/cluster/) with either 100% (no clustering) or 80% homology threshold. The numbers of peptides retained after 80% homology clustering were 16,765 and 23,983 for MHC-I and MHC-II, respectively. (B) Immunogenicity scores generated for various subsets of our epitope datasets. Machine learning was conducted with 32 and 27 the most predictive features for MHC-I and MHC-II, respectively, but without predicted MHC binding information. Evidence: IEDB-derived peptides with functional T cell assay evidence (*i.e.*, without non-IEDB data); Human: epitopes tested in human (*i.e.*, without transgenic animal hosts); Ex vivo: IEDB-derived peptides annotated from direct *exvivo* assays using autologous effector and antigen-presenting cells in human. The numbers of peptides included after applying each filter were 21,162, 13,058, 11,084, and 6,048 for MHC-I, and 31,693, 31,693, 30,188, and 2,023 for MHC-II, respectively. (C) Immunogenicity scores of human epitopes and extrapolated scores for primate and rodent epitopes (N=411 and 8,756 for MHC-I and 76 and 8,445 for MHC-II, respectively). (D and E) Distributions of (D) annotated immunogenicity and (E) immunogenicity scores for the human thymus MHC-I peptidome (Adamopoulou et al., 2013). (F and G) Immunogenicity scores among IEDB-derived (F) MHC-I- and (G) MHC-II-restricted peptides of various origins. HS, *homo sapiens*; HIV, human immunodeficiency virus type I; HBV, hepatitis B virus; HCV, hepatitis C virus; IAV, influenza A virus; DGV, dengue virus; PF, *Plasmodium falciparum*, TB, *Mycobacterium tuberculosis.* (H) Immunogenicity scores among MHC-I-restricted peptides from various sources. HIV, Los Alamos HIV Epitope Database; HCV, Los Alamos HCV Epitope Database. (I) Immunogenicity scores of MHC-I-restricted neoepitopes. NE, neoepitopes from TANTIGEN database. (J) Distributions of immunogenicity scores for neoepitopes grouped by their source antigen proteins. Red points and bars represent median and interquartile ranges, respectively. In (A-I), orange and blue represent immunogenic and non-immunogenic peptides, respectively. In (A-D) and (F-H), bars represent medians. In (E-I), ns, not significant; *, *P* < 0.05; ***, *P* < 0.001; ****, *P* < 0.0001. See also Figure S2, S3, S4, and S5, Data S1, S3, and S4.

We next asked the biological implications of the immunogenicity score framework. First, thymically expressed self-epitopes identified previously (Adamopoulou et al., 2013) were more likely annotated as non-immunogenic (Figure 4D). Moreover, even peptides annotated as immunogenic had a score distribution lower than non-thymus immunogenic and comparable with thymus non-immunogenic counterparts (Figure 4E). Immunogenic peptides from various organisms generally had higher scores than non-immunogenic counterparts, with some exceptions such as MHC-I-restricted peptides derived from human and *Mycobacterium tuberculosis*. Meanwhile, immunogenicity scores were particularly well-suited for HIV and HCV epitopes (Figures 4F and G), and this was also the case in the epitopes from the Los Alamos HIV and HCV immune epitope databases (Figure 4H). Similarly, MHC-I-restricted neoepitopes obtained from TANTIGEN database also exhibited higher scores than non-immunogenic peptides from various origins (Figure 4I). Tumor-associated antigens yielding peptides of high predicted immunogenicity included CTG1B and MAGB1 (Figure 4J).

### The landscape of immunogenicity dynamics by single amino acid mutations

Single amino acid mutations can affect peptide immunogenicity bidirectionally, namely, acquisition and loss of immunogenicity, in sequence space (Figure 5A). We termed the discordance of annotated immunogenicity between neighbors, *i.e.*, single-aa mutants, as “immune transition,” and defined peptides with evidence of immune transition as “transitional.” Note that we cannot define “non-transitional” peptides from our datasets as the lack of evidence of transition may simply result from the lack of assays. We identified 1,360 and 976 transitional peptides for MHC-I and MHC-II, respectively. We noted that immunogenicity scores were less applicable to transitional peptides (Figures 5B and C, and Figure S6), which may result from suboptimal machine learning due to the dearth of transitional peptide data. We next asked whether the integration of neighbor information improves classification. To test this, we constructed neighbor networks for transitional peptides using all possible single-aa mutants (N=232,723 and 292,437 for MHC-I and MHC-II, respectively) simulated computationally, the process we termed “*in silico* mutagenesis.” As expected, the averaged immunogenicity scores classified the immunogenicity of transitional peptides more accurately (Figures 5B and C). To further characterize the position- and residue-specific mutational impacts, we next focused on the immunogenicity dynamics between neighbor pairs. We screened our epitope datasets to identify 6,179 and 3,076 MHC-I and MHC-II-restricted single-aa mutated peptide pairs. Immunogenicity scores changed more dynamically in transitional pairs (Figure 5D). This observation was the case even when focusing on neighbor pairs of common organism origins (Figure 5E). We also noted that score changes between MHC-I neighbor pairs with mutations in their anchor residues were significantly more eminent (Figure 5F). This is interesting as our framework does not utilize any positionally defined features. To gain insights into the residue-specific mutational impacts, we analyzed 2,580,890 and 3,101,092 neighbor pairs from all simulated single-aa mutants of transitional MHC-I- and MHC-II-restricted peptides, respectively. We utilized simulated peptide data because observed neighbor pairs were too sparsely distributed in terms of residue combinations. Heatmap clustering analysis revealed residue-specific impacts on immunogenicity that are partially interpretable with known physicochemical properties [e.g., hydrophobic/aliphatic (V, L, and I), negative (D and E), and aromatic (F and W)]. However, some exceptions were also notable [*e.g.*, positive (K and R) in MHC-I]. Collectively, our systematic characterization delineates the landscape of bidirectional effects of single-aa mutations on immunogenicity in a position- and residue-dependent context.

**Figure 5.**
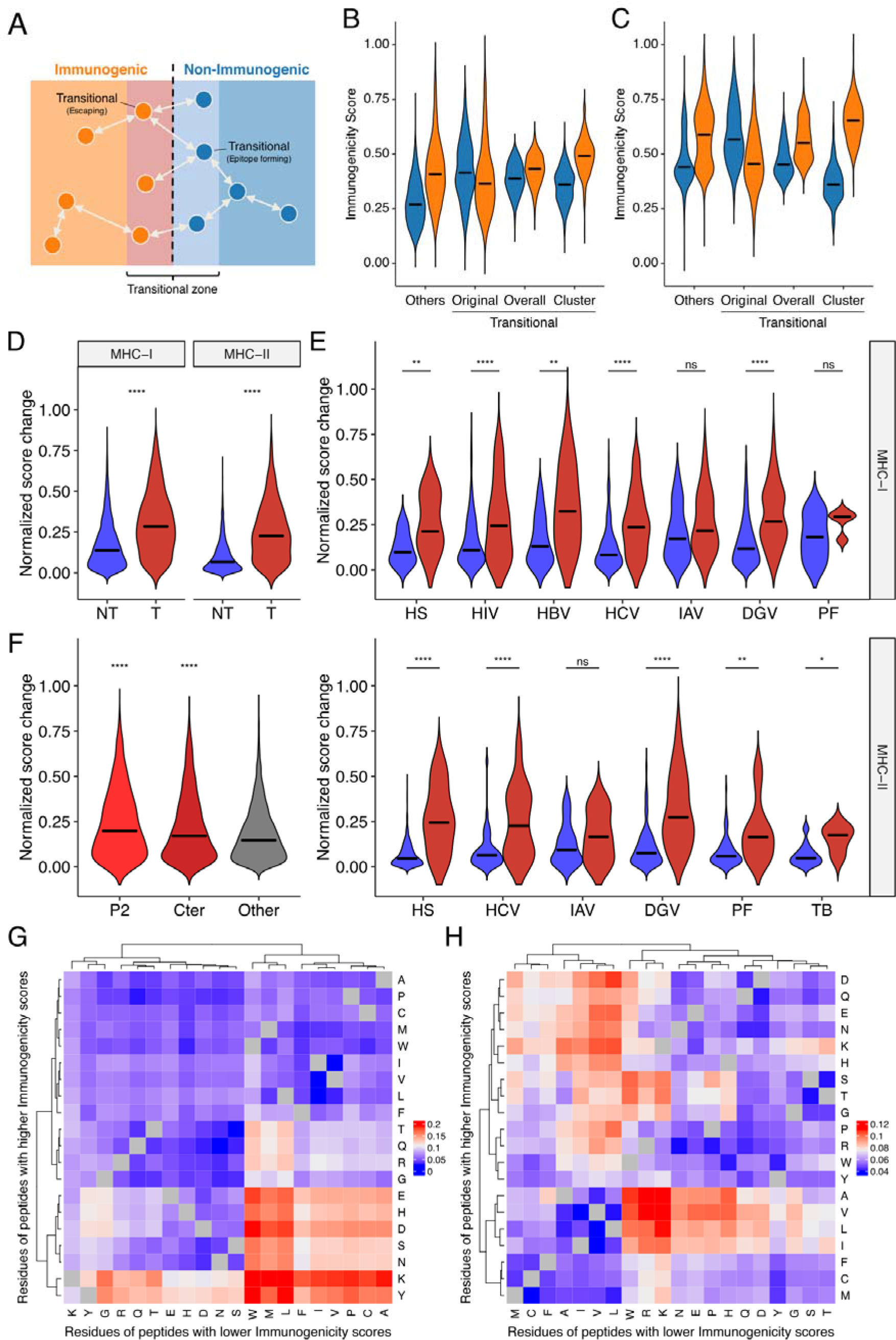
Systematic characterization of the impacts of single amino acid mutations on immunogenicity in sequence space. (A) Schematic of the concept of immune transition in sequence space. We termed the change in immunogenicity between neighbors, *i.e.*, single-aa mutants, as “immune transition,” and defined peptides with evidence of immune transition as “transitional.” (B and C) Immunogenicity scores among (B) MHC-I- and (C) MHC-II-restricted transitional peptides, *i.e.*, peptides with at least one neighbor of opposite immunogenicity annotation in our dataset. We identified 1,360 and 976 transitional peptides for MHC-I and MHC-II, respectively. We expanded their neighbor networks by computing immunogenicity scores for 232,723 and 292,437 all possible single-aa mutants of transitional MHC-I- and MHC- II-restricted peptides, respectively. Original, original immunogenicity scores; Overall, mean immunogenicity scores of all neighbors. Cluster, mean immunogenicity scores of neighbors assigned to the cluster containing the parent peptide. Others, original immunogenicity scores for peptides with no evidence of immune transition (shown for comparison purpose). Orange and blue denote immunogenic and non-immunogenic peptides, respectively. (D-F) Immunogenicity score dynamics among observed neighbor pairs. We identified 6,179 and 3,076 single-aa mutated peptide pairs for MHC-I and MHC-II, respectively. (D) Score dynamics for all possible peptide pairs, regardless of their origins. (E) Score dynamics for pairs of peptides derived from the same organism. Blue and red denote non-transitional and transitional pairs, respectively. (F) Score dynamics of MHC-I peptide pairs stratified by their mutated positions. Cter, C-terminal residue. (G and H) Heatmap clustering analysis of immunogenicity score dynamics by mutating residue pairs. We generated 2,580,890 and 3,101,092 neighbor pairs from 232,723 and 292,437 all possible single-aa mutants of 1,360 and 976 transitional MHC-I- and MHC-II-restricted peptides, respectively. Color represents median normalized score changes. Grey means unobserved residue pairs. In (D-F), ns, not significant; *, *P* < 0.05; **, *P* < 0.01; ****, *P* < 0.0001. See also Figure S6 and Data S5.

### Inference of escaping patterns from neighbor network architecture

Escaping mutations are the principal obstacle for vaccine development. Neither reactivity of the specific T cell clones tested *in vitro* against target epitopes or immune activation after administration *in vivo* does not guarantee a sustainable immune response in the real-world setting because of the possibility of deleterious escaping mutations. We tackled this fundamental problem by quantifying position-specific “escaping potentials” from neighbor network architecture. Figure 6A shows two representative neighbor networks constructed from observed neighbors of immunodominant influenza A virus epitopes GILGFVFTL (GIL) and SRYWAIRTR (SRY). Apparently, GIL seems more robustly immunogenic than SRY because the Cluster4 in the SRY network was enriched with non-immunogenic peptides, suggesting a path for escaping. *In silico* mutagenesis followed by neighbor network clustering analysis revealed position-specific dynamics of predicted immunogenicity (Figure 6B). P6 and P4 were indicated to have the most substantial impact on escaping for GIL and SRY, respectively. Intriguingly, P6 of GIL peptide has been shown to be involved in two hydrogen bonds with two distinct TCR clones F50 and JM22 (Yang et al., 2017). Furthermore, five P4 mutants of SRY peptide has been shown to result in an undetectable level of cytolysis by at least three CTL clones (Bowness et al., 1994). Encouraged by these observations, we next examined two MHC-II-restricted epitopes. GAGSLQPLALEGSLQKRG (GAG) is a self-epitope derived from insulin. Our simulation indicated that P8-11 and P15-16 were required for its immunogenicity. Strikingly, immunodominant epitope LALEGSLQK has been localized previously, showing a complete match with the prediction (Congia et al., 1998). It is worth mentioning that this LALEGSLQK is usually degraded proteolytically during the maturation of insulin molecule. As our framework does not integrate factors affecting intracellular peptide processing and presentation, appropriate peptide presentation onto MHC should be considered a prerequisite for performing immunogenicity score-based analysis. PKGQTGEPGIAGFKGEQGPK (PKG) is another self-epitope derived from collagen. Our simulation indicated that P10, P11, P13, and less evidently, P14 and P16 were required for its immunogenicity. Indeed, mutations at P10, P13, and P16 have been shown to almost diminish the T cell reactivity collected from pre-immunized transgenic mice, and mutations at P14 partially impaired the response (Sakurai et al., 2006). To test if these observations originate from overfitting to target peptides, we repeated the same analysis using probabilistic estimates obtained from machine learning after removing both the target peptide and all of its neighbors. The most prominent positions for every peptide, namely, P6 of GIL, P4 of SRY, P8-P14 of GAG, and P10/P13 of PKG, were reproducibly detected (Figure S7). In the meantime, some of the positions such as P7 of SRY and P16 of PKG became statistically insignificant, indicating that, as well as pan-epitope rules governing immunogenicity, epitope-specific characteristics can also be incorporated when neighbor information is available.

**Figure 6.**
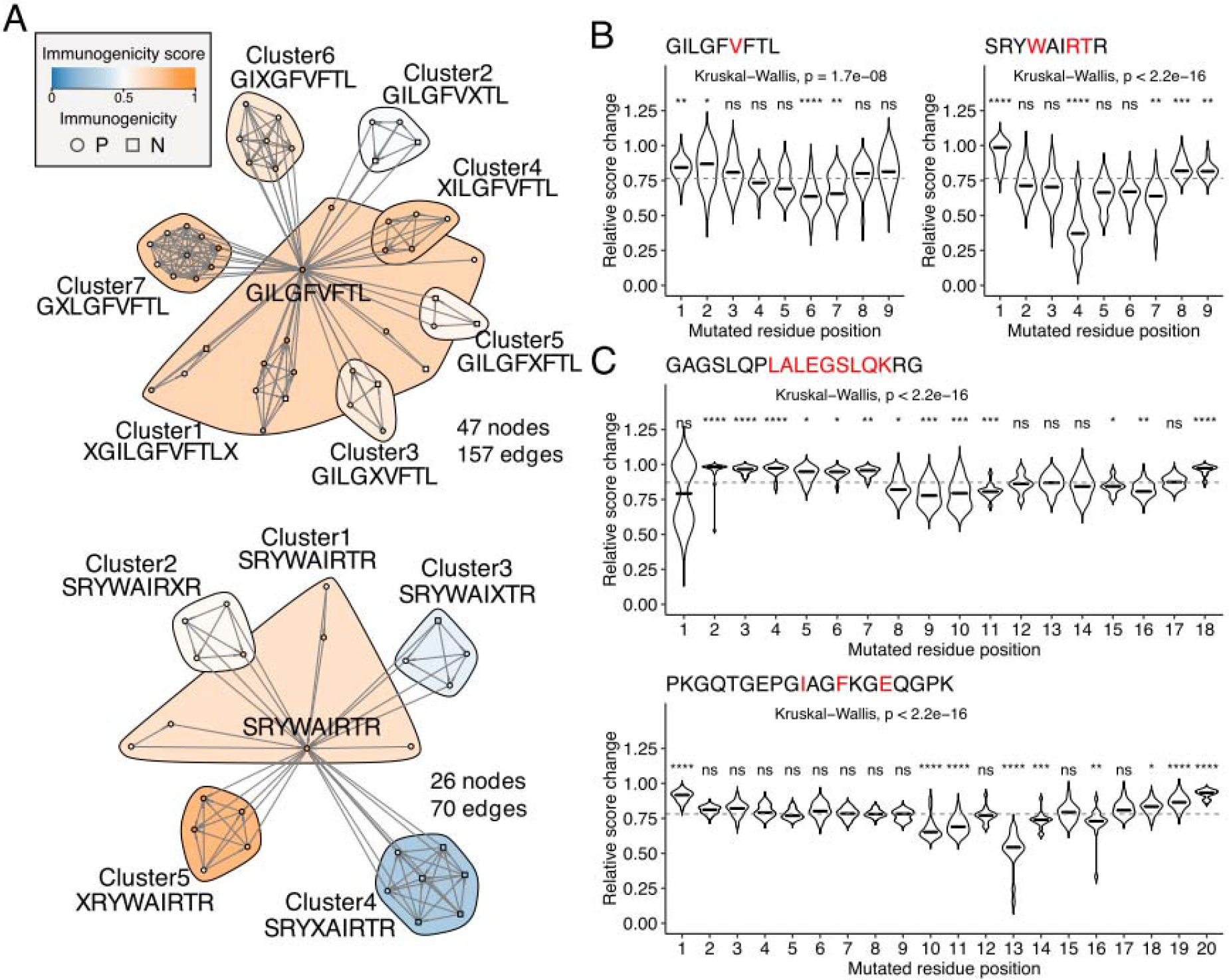
Identification of escape-prone positions from *in silico* mutagenesis and neighbor network analysis. (A) Example neighbor networks of MHC-I-restricted influenza A virus epitopes GILGFVFTL and SRYWAIRTR and their observed neighbors. Consensus sequences are indicated below the cluster IDs. (B and C) Relative score changes estimated from all possible simulated neighbors for (B) MHC-I-restricted influenza A virus epitopes GILGFVFTL and SRYWAIRTR and (C) MHC-II-restricted insulin-derived epitope GAGSLQPLALEGSLQKRG and collagen-derived epitope PKGQTGEPGIAGFKGEQGPK. Red characters indicate residues with known escaping mutations shown by *in vitro* T cell assays. Example peptides were chosenbased on their high immunogenicity scores with manual literature inspection. ns, not significant; *, *P* < 0.05; **, *P* < 0.01; ***, *P* < 0.001; ****, *P* < 0.0001. See also Figure S7.

It is of vast pragmatical significance to invent methodology to computationally prioritize epitopes of high immunogenic potential and low escaping potential. To this end, we defined “escaping potential” as the maximal reduction of cluster-mean immunogenicity score in the neighbor network. Immunogenicity score-escaping potential (IS-EP) plots indicated that, although there was a moderate correlation between IS and EP, considerable variation also existed (Figures 7A and B). Of note, HIV and dengue virus (DGV)-derived epitopes had higher EP than human epitopes, and this trend was not observed in non-immunogenic peptides (Figure 7C). Two representative HIV-derived MHC-I epitopes GGKKKYKL and ITTESIVIW were shown in Figure 7D. As indicated in our analysis, there are known naturally occurring escaping mutations at P3, P5, and P7 of GGKKKYKL (Reid et al., 1996). Although we could not find reliable information on escaping variants of ITTESIVIW, this epitope represents the most common sequence found in the circulating HIV-1 clade B population in the Los Alamos HIV Sequence Database (Du et al., 2016). Likewise, two representative MHC-II epitopes derived from *P. falciparum* and M *tuberculosis* were shown in Figures 7E and F. Indeed, the prevalence of T cell reactivity to the low-EP peptide GKLLSTGLVQNFPNTIISK in Kenyan volunteers was reported to be 25% (Udhayakumar et al., 1995), and, although not precisely the same, 29.4% of South African patients with either latent or active tuberculosis infection were shown to be positive in the ELISpot assay against the epitope VRAVAESHGVAAVLFAATAA (Horvati et al., 2016). Collectively, these observations indicate that our framework enables prioritization of promising epitopes based on their high immunogenic and low escaping potentials *in silico*.

**Figure 7.**
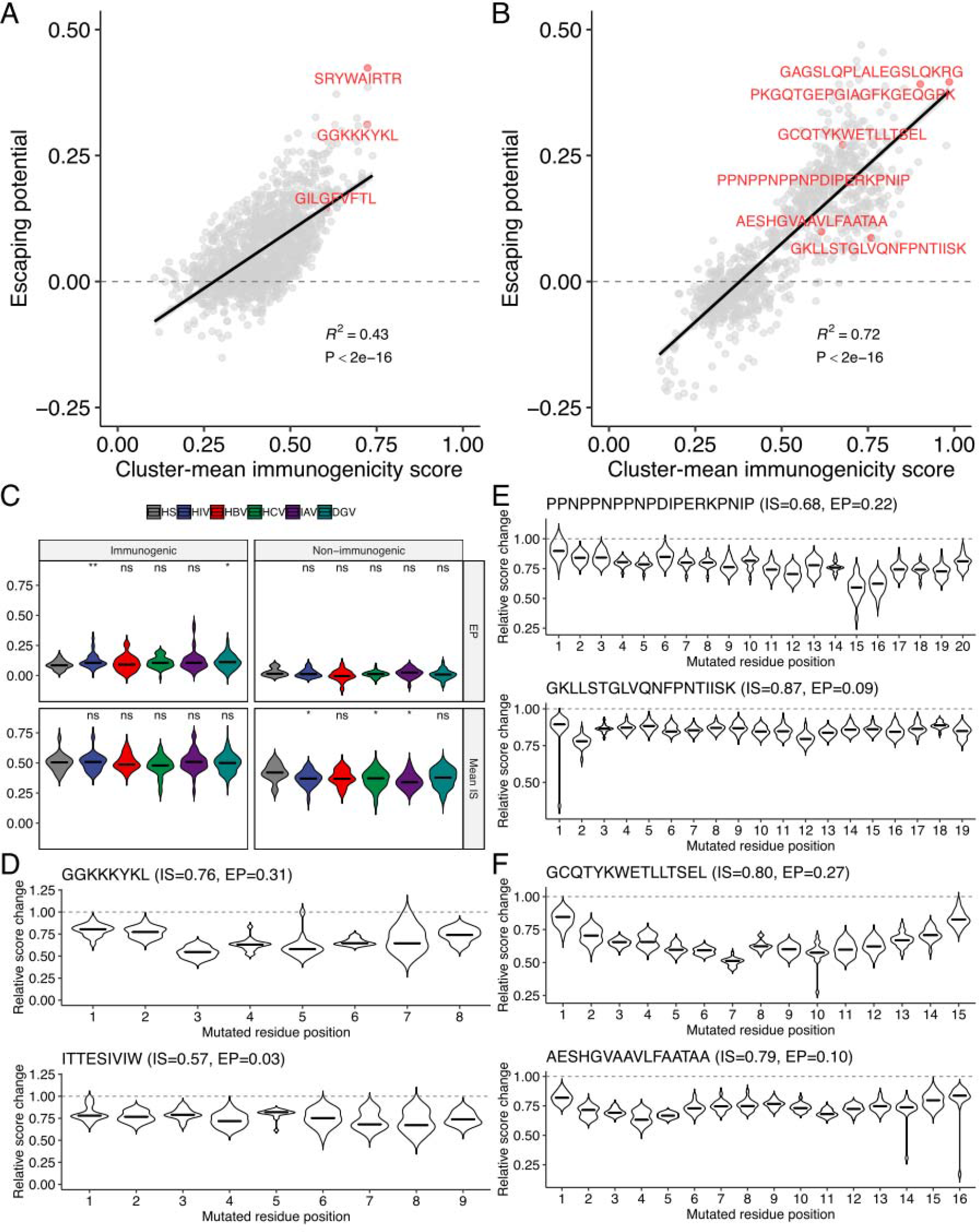
Two-dimensional assessment of epitope quality using the immunogenicity score and escaping potential. (A and B) Immunogenicity score-escaping potential (IS-EP) plots for MHC-I and (B) MHC-II peptides. Cluster-mean immunogenicity scores determined from neighbor network analysis of all simulated neighbors were utilized. Labeled are representative epitopes shown in Figures 6 and 7. (C) IS and EP distributions among pathogen-derived peptides. ns, not significant; *, *P* < 0.05; **, *P* < 0.01. (D-F) Relative score changes estimated from all possible simulated neighbors for (D) representative MHC-I-restricted HIV epitopes, and (E and F) representative MHC-II-restricted epitopes derived from (E) *P. falciparum* and (F) *M. tuberculosis*. In (D-F), representative peptides were chosen based on their high and low EPs with manual literature inspection.

## Discussions

This study provides a proof-of-concept that population-wide aspect of T cell immunity is predictable through probabilistic approach using features recapitulating physiological contacts between MHC-presented peptides and TCR repertoire. It is of note that short (*i.e.*, 3-aa and 4-aa) TCR fragments in combination with a restricted set of AAIndex scales generate features contributing to both MHC-I and MHC-II immunogenicity prediction and also correlating with individual TCR-pMHC affinities, because these commonly utilized parameters might represent MHC-independent pan-epitope principles governing TCR-pMHC interactions. The utilization of short fragment features is consistent with the induced fit model that only a limited complimentary region of TCR is involved at least in initial engagement (Boniface et al., 1999; Garcia et al., 1998; Lee et al., 2004). Meanwhile, the best AAIndex scale BETM990101 reflects interaction energies between amino acid residues under physiological conditions (Betancourt and Thirumalai, 1999), and hence the inverted version of BETM990101, giving larger values for stronger interresidue contacts, is a reasonable scale for contact potential estimation. Indeed, a positive skewness in the contact potential distribution, implying the coexistence of a small and a large proportion of TCR fragments with high and low contact potentials, respectively, was associated with stronger affinities in our analysis. TCR affinity has been correlated with T cell activation (Sykulev et al., 1994), although there remains a debate as to whether other parameters such as avidity, on- and off-rates are also essential (Stone et al., 2009). Further studies are required to fully elucidate the biological relevance of simulated TCR-peptide contact features quintessential for immunogenicity prediction to the thermodynamic parameters of TCR-pMHC interactions and subsequent T cell activation.

Our findings bolster the hypothesis that population-level epitope immunogenicity could indeed be probabilistically determined. Indeed, the concept of immunodominance bases this hypothesis (Yewdell, 2006). For instance, only 0.03% of potential MHC-I-presented peptides derived from vaccinia virus account for >90% of the CD8+ T cell response in B6 mice (Moutaftsi et al., 2006). Similarly, diverse TCR repertoires specific for two immunodominant viral epitopes have been reported in human (Chen et al., 2017). Population-level immunity against various immunodominant epitopes is another line of evidence (Horvati et al., 2016; Udhayakumar et al., 1995). The fact that diverse TCRs can be utilized to achieve protective immunity against immunodominant epitopes in multiple individuals indicates some underlying qualities of epitopes guiding both genomic and thymic “convergent evolution” of human TCR repertoire. To elucidate the hitherto unknown pan-epitope principles governing productive TCR-peptide contacts, we took an inductive approach using a large set of *in vitro* T cell epitope assay results which may vary depending on the specific T cell clones or TCR repertoires utilized. Our findings indicate that a limited set of TCR-peptide contact features reproducibly serve as principal determinants of T cell reactivity. Moreover, we show that the averaged probabilistic estimates, termed immunogenicity scores, are consistently higher in epitopes with positive results from various functional assays. Interestingly, peptides physiologically presented on the thymus have lower immunogenicity scores than non-thymus counterparts, which is consistent with the biased formation of human TCR repertoire owing to thymic negative selection (Adamopoulou et al., 2013; Blackman et al., 1990). Practically speaking, the probabilistic scoring system would be particularly useful for prioritizing peptide candidates to expedite the development of population-level immunotherapies for various infectious diseases and cancer. However, it is notable that immunogenicity scores do not distinguish immunogenic and non-immunogenic peptides from *M. tuberculosis*. It has been shown that T cell epitopes of TB are evolutionary hyperconserved and the bacteria seems to actually benefit from recognition by human T cells (Comas et al., 2010; Coscolla et al., 2015). Further characterization of T cell reactivity for TB epitopes with high and low immunogenicity scores may unveil critical insights into the uniqueness of anti-TB T cell immunity.

One remarkable feature of our immunogenicity score system is that the score dynamics between single-aa mutants are, to some extent, predictive of the mutational impacts on immunogenicity. Moreover, the newly introduce metric termed “escaping potential” allows researchers to quantitatively assess the “robustness” of immunogenicity for any epitope. Practically, it could enable *in silico* epitope screening on the basis of their immunogenicity scores and escaping potentials for developing off-the-shelf targeted immunotherapies available to the general public. In the meantime, it could provide mechanistic insights into the co-evolutionary trajectory of pathogens and general human ancestry. A caveat is that our current model may be suboptimal because only a small fraction of possible sequence space has been experimentally covered so far. Ideally, multiple assays using T cell clones of different individuals should be performed for any peptide to evaluate inter-individual variations. Nonetheless, it is encouraging that our current framework still infers escaping even without using any of the peptide in question and its single-aa mutants during machine learning. Further elucidation of the pan-epitope basis of population-level human T cell immunity has significant clinical and biological implications.

## Materials and Methods

### Experimental design

The purpose of this retrospective study was to identify the essential biochemically relevant features predictive of T cell epitope immunogenicity and to develop a linear coordinate system reflecting the probabilities that those peptides are immunogenic. To this end, public MHC-I and MHC-II-restricted peptide sequence datasets were analyzed through a newly designed computational framework that mimics the molecular scanning of MHC-loaded peptides by human public TCR clonotypes. All analysis is exploratory; no predetermined protocols were applied before initiating the study. All data inclusion and exclusion criteria were described in the following sections. Accounting for missing data values including imputation is not applicable.

### Computational analysis

All computational analyses were conducted using R ver. 3.5.0 (https://www.r-project.org/) (R Core Team, 2018). The latest versions of R packages were consistently used. Compiled datasets and essential in-house functions are available as the R package *Repitope* on GitHub (https://github.com/masato-ogishi/Repitope). Other scripts are available upon request.

### Peptide sequence datasets

HLA-I-restricted peptide sequences of 8-aa to 11-aa lengths with T cell assay results were collected from public databases [Immune Epitope Database (IEDB, as of May 7^th^, 2018) (Fleri et al., 2017), the best-characterized CTL epitopes from Los Alamos National Laboratory (LANL) HIV Sequence Database (Llano et al., 2013), LANL HCV Sequence Database (Kuiken et al., 2005), EPIMHC (Reche et al., 2005), MHCBN (Lata et al., 2009), and TANTIGEN (Olsen et al., 2017)] and previous publications (Calis et al., 2013; Chowell et al., 2015; Tung et al., 2011). HLA-II-restricted peptide sequences of 11-aa to 30-aa lengths with T cell assay results were collected from the IEDB database (as of May 7^th^, 2018). As for peptides retrieved from the IEDB database, only those with evidence of functional T cell response were included. The list of evidence included is as follows: activation, antibody help, CCL2/MCP-1 release, CCL3/MIP-1a release, CCL4/MIP-1b release, CCL5/RANTES release, CXCL10/IP-10 release, CXCL9/MIG release, cytotoxicity, decreased disease, degranulation, disease exacerbation, GM-CSF release, granulysin release, granzyme A release, granzyme B release, IFNg release, IL-10 release, IL-12 release, IL-13 release, IL-17 release, IL-17A release, IL-1b release, IL-2 release, IL-21 release, IL-22 release, IL-23 release, IL-3 release, IL-4 release, IL-5 release, IL-6 release, IL-8 release, lymphotoxin A/TNFb release, pathogen burden after challenge, perforin release, proliferation, protection from challenge, survival from challenge, T cell-APC binding, T cell help, TGFb release, TNF release, TNFa release, tolerance, tumor burden after challenge, and type IV hypersensitivity (DTH). Peptides presented on non-human MHC molecules were discarded, whereas those presented on HLA molecules in non-human hosts (*e.g*., transgenic mice) were included. In this manner, 21,162 and 31,693 HLA-I and HLA-II-restricted peptides were identified, respectively, of which 1,873 (8.9%) and 4,505 (14.2%) had contradicting annotations on immunogenicity. In such cases, peptides with at least one positive T cell assay result were considered immunogenic to maximize the breadth of population-wide immunity included in the analysis. Eventually, 6,957 (32.9%) and 16,642 (52.5%) HLA-I and HLA-II-restricted peptides were considered immunogenic, respectively. Collected peptide sequences and annotations are summarized in Data S1.

Sequences of peptides restricted on non-human MHC molecules were collected from the IEDB database (as of as of May 7^th^, 2018) as described above. The following species were considered primate: bonobo, chimpanzee, gorilla, marmoset, and rhesus macaque. Meanwhile, mouse and rat were considered rodent. Eventually, 411 and 8,756 MHC-I-restricted peptides, and 76 and 8,445 MHC-II-restricted peptides were identified for primates and rodents, respectively. Identified epitopes and annotated information can be found in Data S1.

### TCR sequence datasets

TCR repertoire datasets were collected from NCBI Sequence Read Archive (SRA) and a previous study led by Britanova *et al.* (Britanova et al., 2014). Repertoire datasets derived from healthy donors were searched through SRA, and the following BioProjects were included: PRJNA389805, PRJNA329041, PRJNA273698, PRJNA258001, PRJNA229070, PRJNA79707, and PRJNA79435. Also, pooled sequence data from 39 healthy donors were retrieved from the paper by Britanova *et al.* In this manner, a total of 206 datasets, of which only five were pooled datasets, were collected. Pooled and individual datasets were derived from a total of 561 and 103 healthy donors, respectively. Fastq files were obtained using fastq-dump script with the following options: --gzip --skip-technical --readids --read-filter pass --dumpbase --split-files-clip --accession [SRA run number]. A total of 23,006,555 CDR3β sequences were extracted using MiXCR software (Bolotin et al., 2015).

Public TCR clonotypes, the ones commonly observed in multiple datasets, were identified from the pool of CDR3β sequences mentioned above. Britanova *et al.* (Britanova et al., 2014) reported that 10,691 clonotypes were shared between at least six of 39 (15.4%) donor-derived top 100,000 clonotype sets. Based on their findings, we decided to extract CDR3β sequences identified in at least 22 out of 206 (10.8%) different datasets, and we subsequently identified 191,326 (0.83%) public clonotypes. A randomly sampled 10,000 clonotypes were used for TCR-peptide contact potential profiling.

### TCR-peptide contact potential profiling (CPP)

T cell epitopes presented on the MHC molecules must be recognized by the TCRs of CD8+ CTLs and CD4+ Th cells with sufficient affinity to trigger subsequent immunological cascades. Undoubtedly, the vast majority of TCR repertoire is not involved in recognition of any given peptide. Moreover, considering the substantial conformational flexibility observed upon binding of TCRs to pMHC complexes, one can assume that only a subset of residues of the peptide and the epitope-recognition domain, namely, the CDR3β loop, serve as a “seed” of intermolecular docking during the early phase of molecular scanning (Boniface et al., 1999; Bridgeman et al., 2012). With these in mind, identifying the best contact site between a given pair of peptide and CDR3β sequences is conceptually similar to solving a local pairwise sequence alignment problem. One caution is that higher scores must be given to more strongly interacting residue pairs instead of more biochemically similar residue pairs as opposed to ordinary alignments. For this purpose, amino acid pairwise contact potential (AACP) scales from the AAIndex database (Kawashima et al., 2008) (http://www.genome.jp/aaindex/AAindex/list_of_potentials) were adopted to generate custom substitution matrices. As stronger interresidue interactions yield smaller free energy values (negative numbers), and as the Smith-Waterman local alignment algorithm attempts to maximize the alignment score of a given sequence (TCR fragment) against the whole target sequence (peptide), the optimal pairwise alignment using a custom substitution matrix derived from the sign-inverted version of a pairwise contact potential scale would in principle correspond to the best intermolecular contact. For comparative purposes, both inverted and non-inverted versions were tested. Values were rescaled to a range from zero to one for subsequent analyses. A set of non-inverted, non-rescaled AACP scales is provided as Data S6. For repertoire-wide CPP analysis, the 191,326 pooled public CDR3β sequences were randomly down-sampled to 10,000 sequences with the relative abundance ratios retained. The sequences were then fragmented by a sliding window strategy to generate a fragment library. Since the interacting orientations (either forward parallel or antiparallel) are unknown for most of the cases, reversed CDR3β sequences were also fragmented and combined. The sizes of fragments were from 3-aa to 8-aa and from 3-aa to 11-aa for MHC-I and MHC-II predictions, respectively. For single-TCR CPP (sCPP) analysis, every single TCR instead of public TCR repertoire was used to generate a fragment library. To perform sequence alignments, the *pairwiseAlignment* function implemented in the *Biostrings* package in *Bioconductor* (https://www.bioconductor.org/) (Gentleman et al., 2004) was utilized. This function seeks an optimal alignment which maximizes the overall alignment score defined as a sum of pairwise scores. Alignment type was set “global-local” to obtain an optimal alignment of a given set of TCR fragments against the consecutive subsequences of any given peptide. Gaps were not allowed. A set of alignment scores was summarized by calculating representative statistics. Following statistics were calculated using the functions implemented in the *psych* package: mean (Mean), standard deviation (SD), standard error of the mean (SE), median (Med), median absolute deviation (MAD), interquartile range (IQR), 10% quantile (Q10), 90% quantile (Q90), skewness (Skew), and kurtosis (Kurt). Predictive features were generated by combining the fragment length, the AACP scale, and the type of statistics. In this manner, 700 CPP features per one fragment length were generated for each of the peptides.

### Peptide descriptors

Apart from CPP features, sequence-based estimates of physicochemical properties were also calculated. Each peptide sequence was converted into a set of consecutive fragments of a defined amino acid length, and peptide descriptors were calculated against each of the fragments using functions in the *Peptide* package. Following functions were utilized: *aIndex, blosumIndices, boman, charge, crucianProperties, fasgaiVectors, hmoment, hydrophobicity, instaIndex, kideraFactors, mswhimScores, pI, protFP, vhseScales*., and, *zScales*. The distributions of the values were summarized similarly to CPP features. Additionally, 20 categorical features indicating whether the peptide of interest is free from a specific amino acid residue were included. The peptide length was also included as a feature.

### Feature selection

Preprocessing followed by importance-based feature selection was repeated ten times with different random seeds. The analysis workflow consisted of the following steps. First, the peptide set was randomly split in a ratio of 4:1 for the training and testing subdatasets. Second, using the training subdataset, a preprocessing function was defined; numerical features were centered and scaled; a peptide length and categorical features representing existence or absence of each residue remained unchanged. This preprocessing function was also applied to the testing subdataset. Third, highly correlated features were removed, with the threshold of correlation coefficient being 0.75. Fourth, features were filtered based on the importance values calculated using the *generateFilterValuesData* function with the random forest (RF) method *(randomForestSRC.rfsrc)* implemented in the *mlr* package (Bischl et al., 2016). The 100 most important features were retained unless otherwise stated. In the first five repeats, the names of the peptide descriptors and the CPP features were ordered in ascending order. In the latter five repeats, on the other hand, the names of those features were ordered in descending order. In this manner, the inherent bias of feature selection owing to the lexicographical order of the names of features was avoided. Finally, features kept in all of the ten repeats were defined as the most predictive features and were utilized hereafter.

### TCR-pMHC structure analysis

The structures of TCR-pMHC complexes with known experimentally determined affinities were collected manually from the literature using the ATLAS database (https://zlab.umassmed.edu/atlas/web/) (Borrman et al., 2017). A total of 95 structures were retrieved, of which 82 contained non-mutated (wildtype) TCRs. Contact sites were identified using the PRODIGY server (Vangone and Bonvin, 2015; Xue et al., 2016). TCR contact footprints were defined as the distributions of the numbers of contacts at each of the peptide positions.

For the analysis of correlation with affinities, sCPP features were computed as described in the above sections, and the single best feature was selected from a univariate analysis for each of the AAIndex scales. Multivariate regression with stepwise feature selection was conducted using the *stepAIC* function implemented in the *MASS* package. The variance inflation factors (VIFs) were computed using the *vif* function implemented in the *car* package.

Sequence permutation experiments were performed as follows. Peptide sequences were replaced with a set of random sequences of the same lengths. In contrast, because of the fragmentation strategy employed, a simple replacement would lead to incomplete disruption of hidden but essential motifs. Therefore, we instead replaced the TCR-derived fragment library with a randomly chosen set of random fragments that do not overlap with the original fragment library. Permutation experiments were iterated 1000 times.

### HLA binding prediction

Predicted binding strength against twelve representative HLA-I alleles (A*01:01, A*02:01, A*03:01, A*24:02, A*26:01, B*07:02, B*08:01, B*27:05, B*39:01, B*40:01, B*58:01, and B*15:01) from NetMHC 4.0 (Andreatta and Nielsen, 2016) and six representative HLA-II alleles (DRB1*0101, DRB3*0101, DRB4*0101, DRB5*0101, DPA1*0103-DPB1*0101, and DQA1*0101-DQB1*0201) from NetMHCIIpan 3.2 (Jensen et al., 2018) for MHC-I and MHC-II immunogenicity prediction, respectively, were incorporated in the machine learning process for immunogenicity prediction because the stability of the peptide-MHC complex is a known correlate of immunogenicity (Rasmussen et al., 2016). Default parameters were used for prediction. Thresholds of percentile ranks for strong and weak binders were set at 0.5% and 2% in MHC-I, and 2% and 10% in MHC-II, respectively.

### Peptide clustering by the IEDB Cluster Analysis Tool

The IEDB Epitope Cluster Analysis Tool (http://tools.iedb.org/cluster/) was utilized to test the effect of sequence-level peptide homology on immunogenicity prediction. Clustering was performed with the homology threshold of 80%. When clustering filter was applied, only a single peptide was randomly chosen from each of the clusters before feature preprocessing in the machine learning workflow.

### Immunogenicity score

The most predictive peptide descriptors and CPP features, with or without predicted HLA binding-based features, were compressed into a linear coordinate system through machine learning. We chose extremely randomized trees (ERT) algorithm implemented in the *extraTrees* package (Geurts et al., 2006) from preliminary experiments. ERT is a derivative of RF and is more computationally efficient and at the same time less prone to overfitting. The model-specific parameter *mtry* was set to be 5. Other hyperparameters were set as defaults. Class weights were provided to compensate for imbalanced class distributions. Probabilistic estimates were computed by conducting five times repeated five-fold cross-validations (CVs). This strategy ensures that any peptide is subjected to five times repeated predictions by models trained from a set of peptides that do not contain the peptide in question. The averaged probability estimates from machine learning were termed “immunogenicity scores.” Unless otherwise stated, we utilized immunogenicity scores from machine learning without predicted HLA binding-based features. For peptides not included in our epitope dataset, immunogenicity scores are defined as the averaged probability estimates of the 25 ERT models generated during CVs. Prediction variances were evaluated by computing coefficients of variance which are defined as standard deviations divided by means.

### Immunogenicity prediction by the IEDB Class I Immunogenicity Tool

Immunogenicity was predicted on the HLA-I-restricted peptide set using the IEDB Class I Immunogenicity Tool (22) (http://tools.iedb.org/immunogenicity/) with default settings.

### Neighbor network analysis

Single amino acid mutations, namely, substitutions, insertions, and deletions, have multifaceted effects on the immunogenicity of MHC-presented peptides. To systematically analyze their effects on immunogenicity, we adopted a network-style representation of the epitope data structures, termed a “neighbor network,” where pairs of peptide sequences (nodes) with just one edit distance were defined as “neighbors” and regarded as edges. Each edge was directed from the peptide with a lower immunogenicity score to the peptide with a higher score. Edge weight was defined as the ratio of the lower score to the higher score so that less immunogenically similar peptides were mapped more distant from each other in sequence space due to the smaller edge weight. Clustering was performed using a walktrap algorithm implemented in the *igraph* package (Csárdi and Nepusz, 2006). For each cluster, sequences were aligned using the ClustalW algorithm implemented in the *msa* package (Bodenhofer et al., 2015). A consensus sequence was generated using the method implemented in the *Biostrings* package with an ambiguity threshold of 0.5. Gaps were treated equally to ambiguities. Gaps/ambiguities were expressed as “X.”

### *In silico* mutagenesis analysis

For a given “parental” peptide, the neighbor network was computationally expanded by simulating all possible single amino acid substituted mutant peptides. Insertions and deletions were not taken into consideration. These artificially introduced mutations could affect various aspects of T cell immunity, including aberrant proteolytic cleavage and altered MHC presentation, these possibilities were not rigorously examined since our primary interest is on exploring the net immunogenicity of mutated peptides if appropriately presented and scanned by TCRs. Immunogenicity score computation and neighbor network analysis were conducted as described.

### Immune transition

Loss of immunogenicity due to mutation is also called escaping, evasion, and sometimes immunoediting. In the present study, we refer to this phenomenon as escaping. In contrast, there is no appropriate terminology for the mutation-driven acquisition of immunogenicity, and thus we refer to this phenomenon as “epitope/neoepitope formation.” For the sake of simplicity, we do not take into consideration epitope-extrinsic mechanisms such as impaired intracellular antigen processing, somatic loss of HLA heterozygosity (Chowell et al., 2018; McGranahan et al., 2017), and checkpoint-mediated T cell exhaustion and apoptosis (Pardoll, 2012) in this work. Escaping and epitope formation due to mutations can be understood as two sides of the same coin in sequence space, and therefore, we propose a higher-order concept, “immune transition,” which is defined as a change in immunogenicity between any MHC-presented peptide and its single amino acid variant. Note that the synergistic effects of two or more mutations on the changes in immunogenicity are beyond the scope of the present study.

To further characterize the theoretical boundary of immunogenic and non-immunogenic peptides, we categorized peptides with evidence of immune transition found in our dataset as “transitional” and analyzed separately. Note that we cannot certainly define “non-transitional” peptides, as the lack of immune transition in our dataset may merely reflect the lack of experimental evaluation. For transitional peptides, *in silico* mutagenesis followed by neighbor network analysis was conducted. Mean immunogenicity scores of both all single amino acid mutants and those within the same cluster to the parental peptide were computed.

### Immunogenicity score dynamics

For quantitative comparison of immunogenicity for a given pair of peptides, either the ratio of immunogenicity scores (divided by the lower score), termed “relative score change,” or the absolute difference of immunogenicity scores divided by the lower score, termed “normalized score change,” was used as a metric. To systematically explore the position-specific mutational impacts for a given parental peptide, relative score change was computed for every pair of computationally simulated mutants. For heatmap analysis, peptide pairs were grouped based on their mutated residues, and median normalized score changes were calculated for each group. Heatmaps were visualized using the *ComplexHeatmap* package (Gu et al., 2016).

### Escaping potential

Escaping potential of a given peptide was defined as the difference between the mean immunogenicity score of the cluster containing the parental peptide and the minimum mean immunogenicity score of the other clusters in the neighbor network containing all possible simulated mutants. A negative escaping potential indicates that the parental peptide resides in the least immunogenic cluster.

### Statistical analysis

Non-parametric hypothesis testing methods (Kruskal-Wallis test and Wilcoxon rank sum test for continuous variables, and Fisher’s exact test for categorical variables) were consistently employed unless otherwise stated. In the case of multiple comparisons, *P*-values were FDR-adjusted according to the method originally proposed by Benjamini and Hochberg (Benjamini and Hochberg, 1995). When reporting measures of central tendency and dispersion, data were presented as medians and interquartile ranges unless otherwise stated. AUC values were calculated based on ten-fold CVs, and their 95% confidence intervals were inferred from influence curves using the method implemented in the *cvAUC* package (Ledell et al., 2015). All statistical analyses were conducted in R (R Core Team, 2018).

## Author contributions

M.O. conceived the idea, designed the study, performed computational analyses, and drafted the manuscript; M.O. and H.Y. wrote the manuscript.

## Acknowledgments

We thank Ryo Ueno for helpful discussions. This work is not supported by any external funding sources.

### Declaration of interest

The authors declare no competing interests.

## Supplementary Materials

### Supplementary Figures

**Figure S1.**
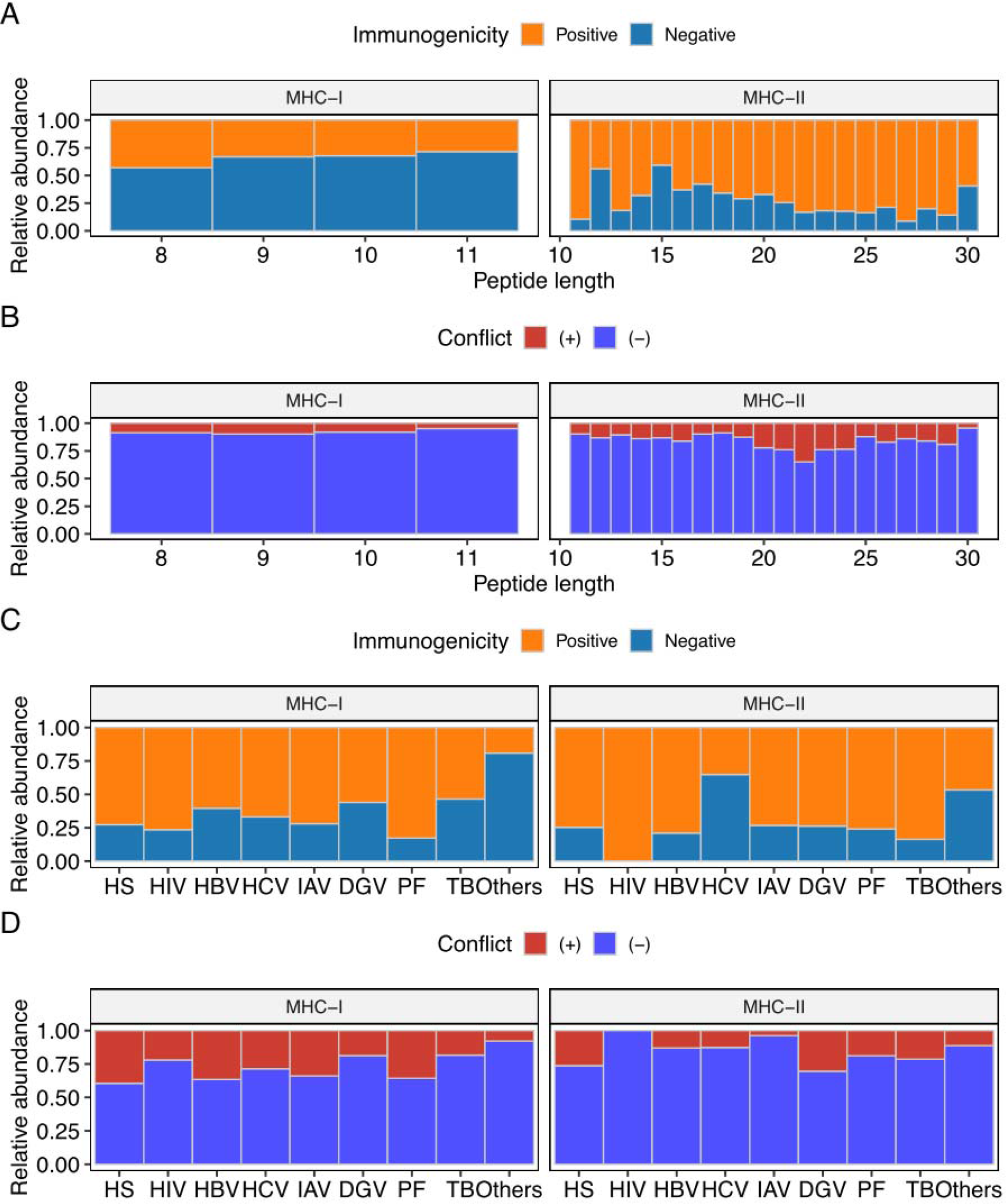
Base characteristics of the epitope dataset, Related to Figure 1. (A and B) Relative abundance of (A) immunogenicity annotations and (B) conflicts in immunogenicity annotations among peptides of different lengths. (C and D) Relative abundance of (C) immunogenicity annotations and (D) conflicts in immunogenicity annotations among peptides of different origins. HS, homo sapiens; HIV, human immunodeficiency virus type 1; HBV, hepatitis B virus; HCV, hepatitis C virus; IAV, influenza A virus; DGV, dengue virus; PF, Plasmodium falciparum; TB, Mycobacterium tuberculosis; Others, peptides derived from other organisms (peptides with no source organism annotation were excluded).

**Figure S2.**
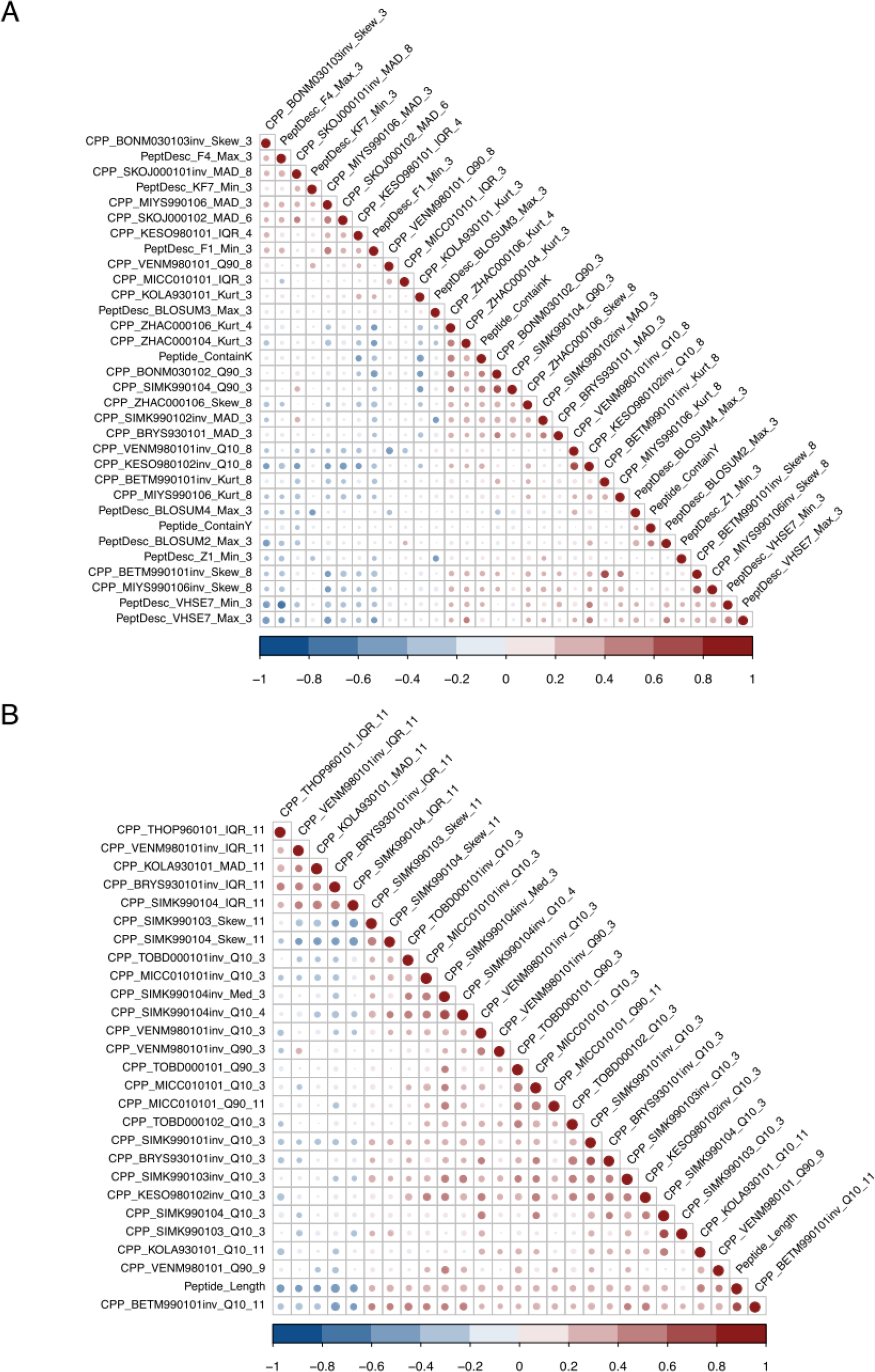
Correlograms of the minimal sets of features predictive of epitope immunogenicity, Related to Figure 3. (A) Features for MHC-I-restricted peptides (N=32). (B) Features for MHC-II-restricted peptides (N=27). Features were reordered by the hierarchical clustering algorithm.

**Figure S3.**
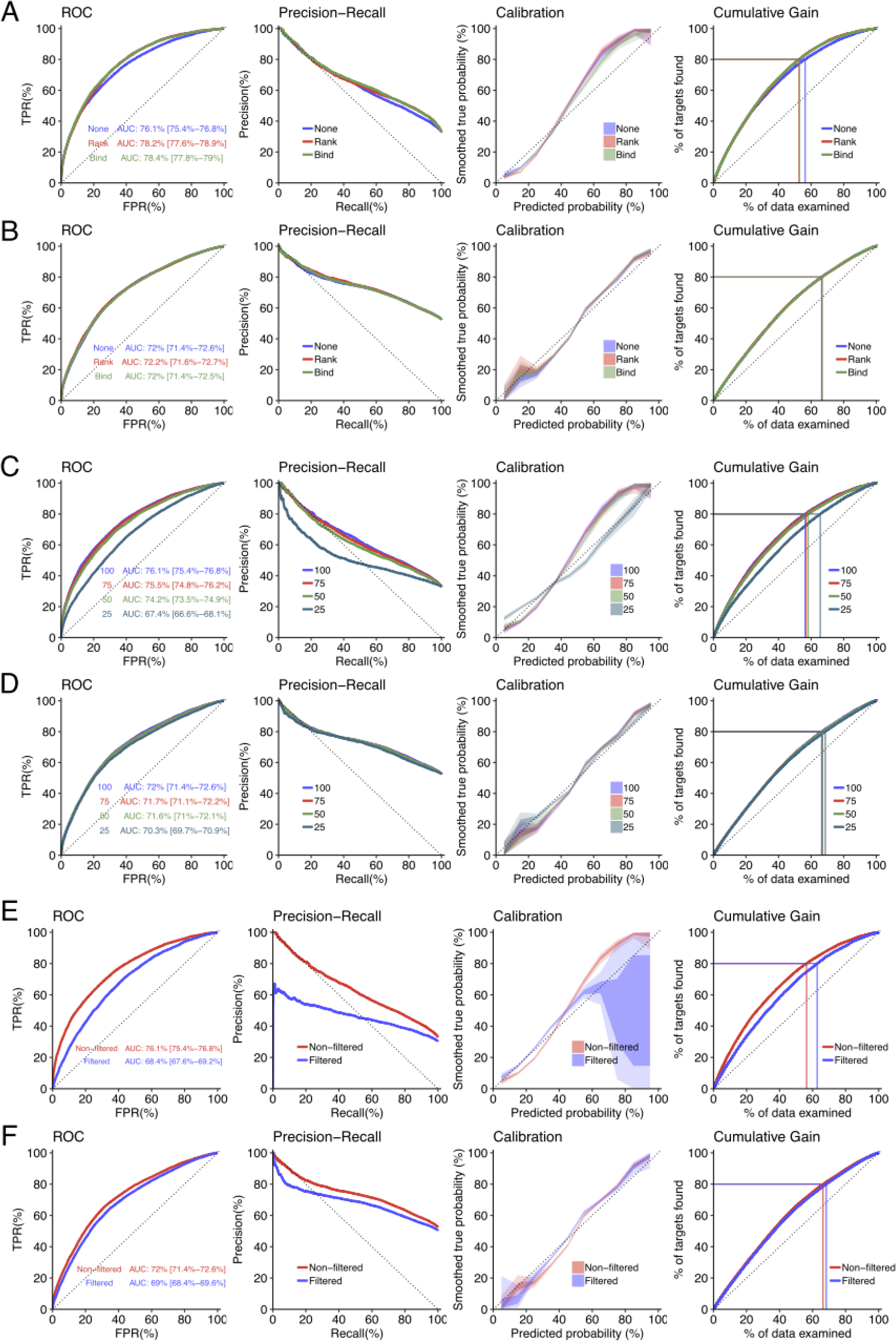
Robustness analysis of immunogenicity predictions, Related to Figure 4. Performances of probabilistic estimates computed through machine learning under various conditions were examined. (A and B) Addition of MHC binding prediction results as features for machine learning. Rank, percentile ranks; Bind, binding strength categories. (A) MHC-I-restricted peptides. (B) MHC-II-restricted peptides. (C and D) Machine learning with smaller cutoff values for feature selection. (C) MHC-I-restricted peptides. The actual numbers of features retained after feature selection were 32, 26, 19, and 7, respectively. (D) MHC-II-restricted peptides. The actual numbers of features retained were 27, 23, 21, and 12, respectively. (E and F) Peptide homology-based clustering before machine learning. The IEDB Epitope Cluster Analysis Tool (http://tools.iedb.org/cluster/) was used to cluster the peptides with the homology threshold of 80%. (E) MHC-I-restricted peptides. A total of 16,765 peptides were retained after clustering. (F) MHC-II-restricted peptides. A total of 23,983 peptides were retained after clustering. Nonfiltered, all peptides. Filtered, single peptide randomly chosen from each of the clusters before preprocessing.

**Figure S4.**
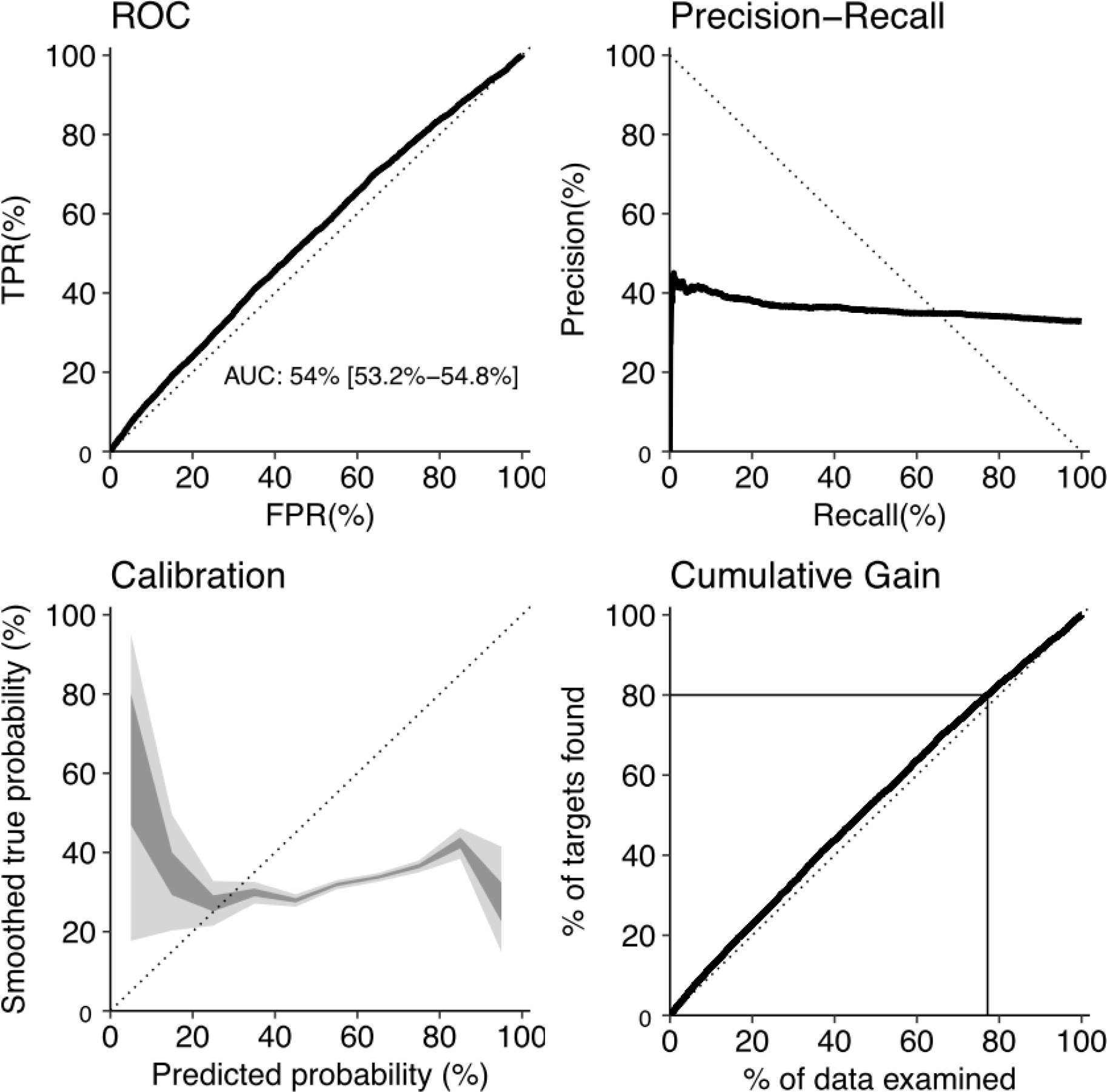
Immunogenicity prediction of HLA-I-restricted peptides by the IEDB Class I Immunogenicity Tool, Related to Figure 4. Classification performances were evaluated using the prediction scores (rescaled to 0-1 range) derived from the IEDB Class I Immunogenicity Tool (Calis et al., 2013) (http://tools.iedb.org/immunogenicity/) with default settings. Note that peptides used for training the model in the original paper were not excluded from prediction, which theoretically overestimates its predictive performance.

**Figure S5.**
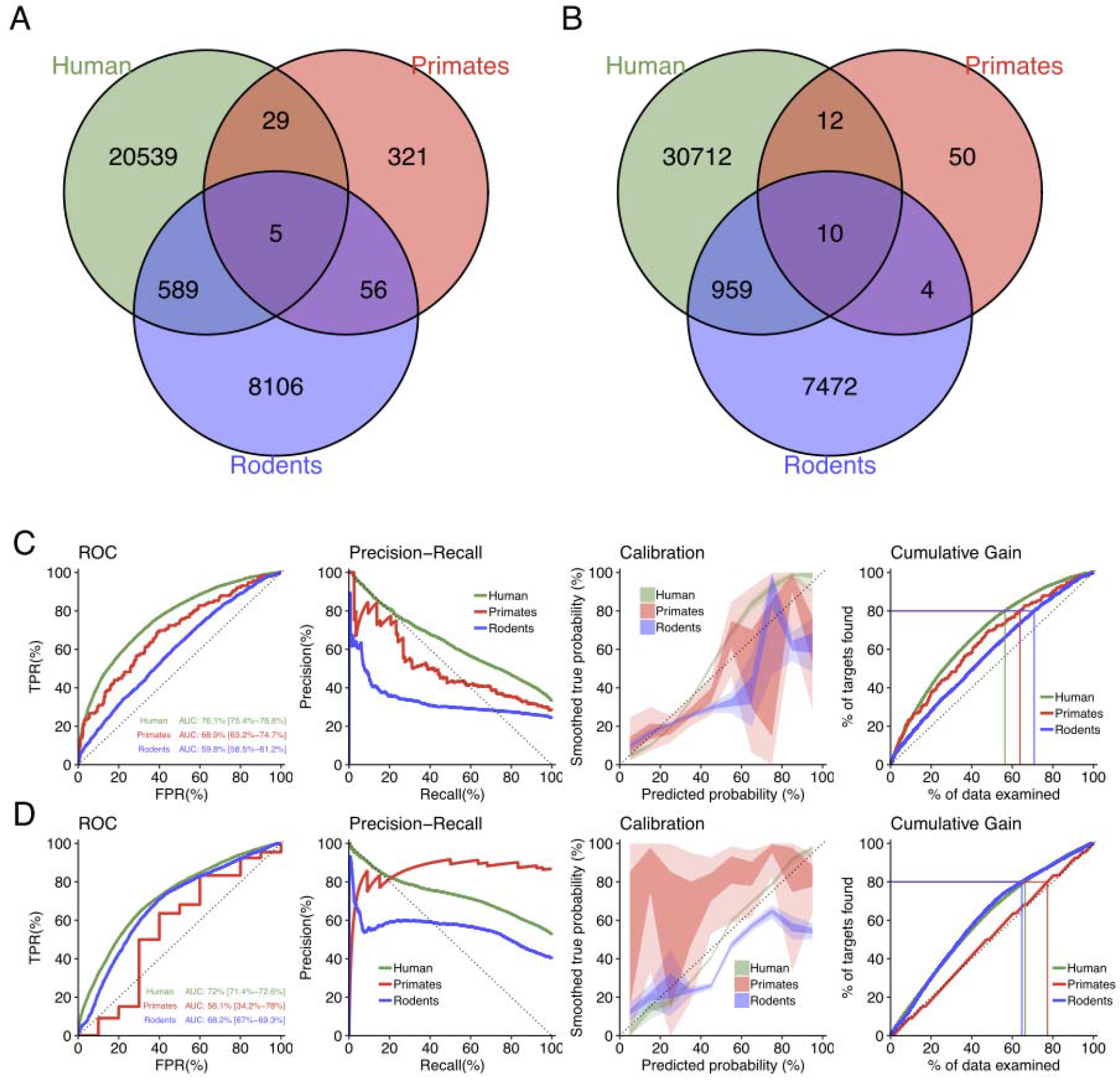
Extrapolation of the human immunogenicity prediction framework for primate and rodent epitopes, Related to Figure 4. (A and B) Overlaps between (A) MHC-I-restricted and (B) MHC-II-restricted peptides presented on MHC molecules of different species. (C and D) Predictive performances of the probabilities of immunogenicity for (C) MHC-I-restricted and (D) MHC-II-restricted peptides. Probabilities for non-human peptides were estimated by applying the classifiers trained on the human peptide datasets.

**Figure S6.**
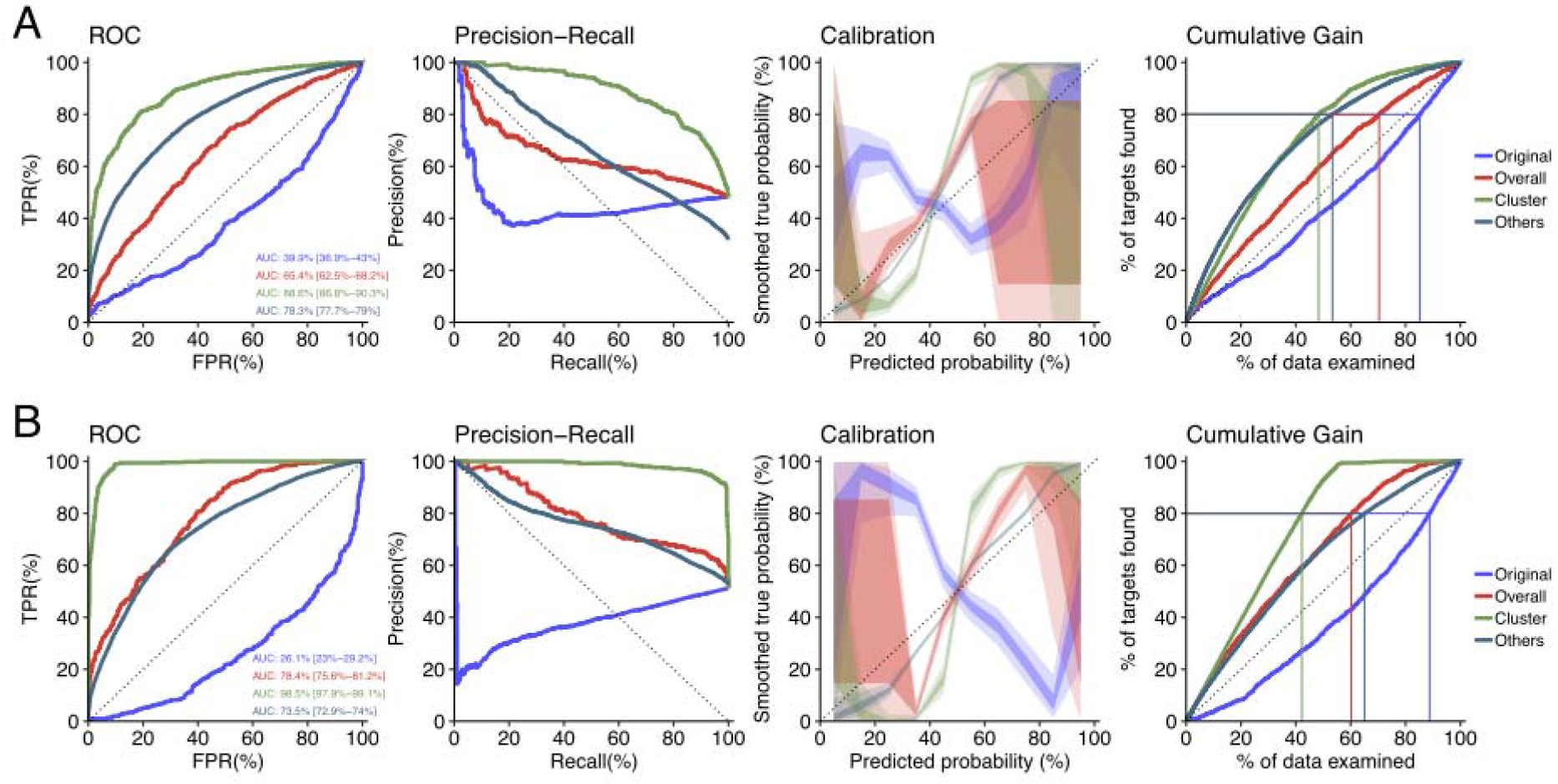
Improvement of predictive performances of immunogenicity scores for transitional peptides by integrating the neighbor network architectures in sequence space, Related to Figure 5. Predictive performances of the immunogenicity scores of (A) MHC-I- and (A) MHC-II-restricted transitional peptides, *i.e*., peptides with at least one neighbor of opposite immunogenicity annotation in our dataset, were evaluated. We identified 1,360 and 976 transitional peptides for MHC-I and MHC-II, respectively. We expanded their neighbor networks by computing immunogenicity scores for 232,723 and 292,437 all possible single-aa mutants of transitional MHC-I- and MHC-II-restricted peptides, respectively. Original, original immunogenicity scores; Overall, mean immunogenicity scores of all neighbors. Cluster, mean immunogenicity scores of neighbors assigned to the cluster containing the parent peptide. Others, original immunogenicity scores for peptides with no evidence of immune transition (shown for comparison purpose).

**Figure S7.**
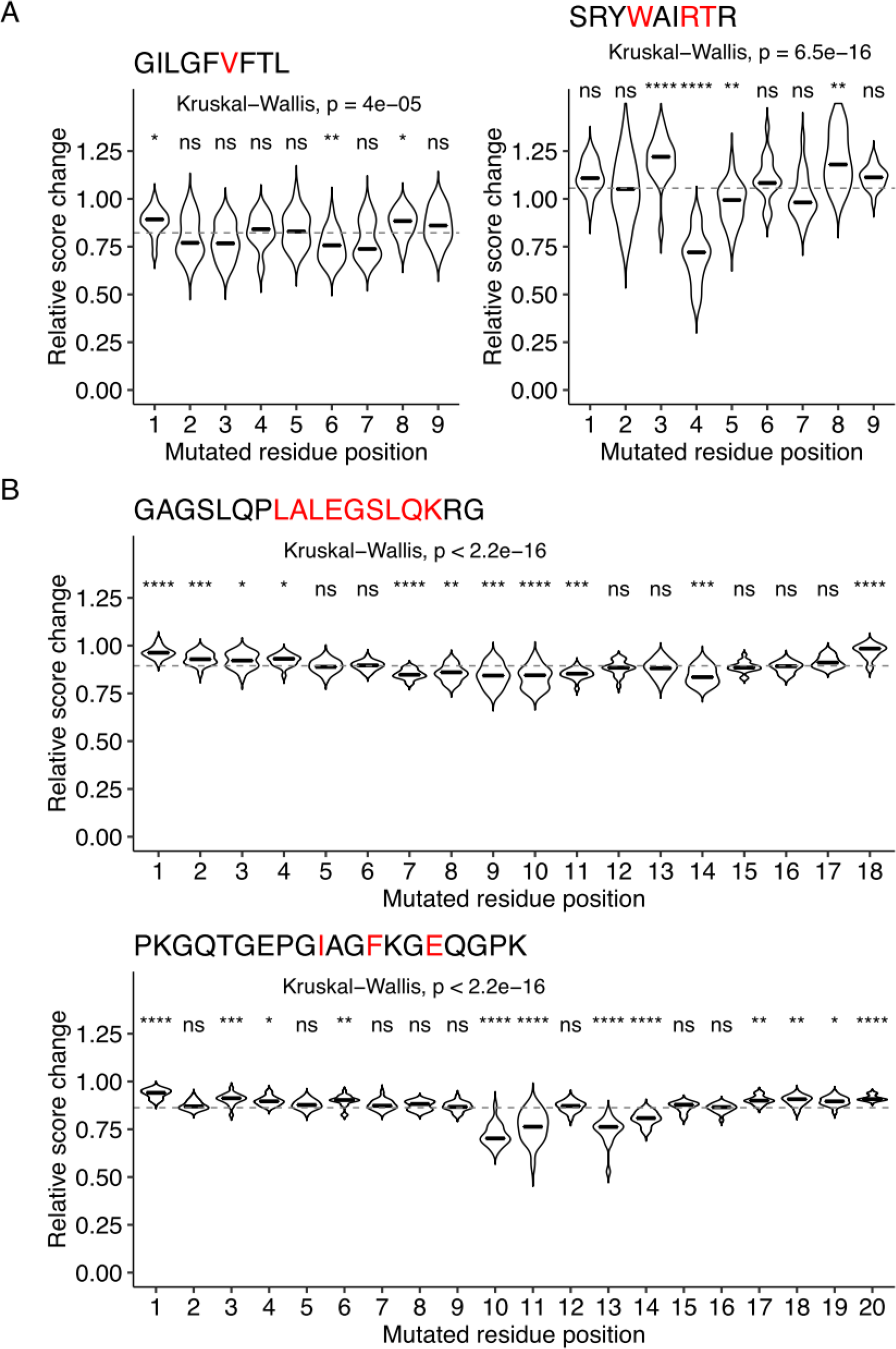
Assessment of position-specific escaping potentials for representative peptides using probabilistic estimates of immunogenicity inferred without target peptides and their neighbors during machine learning, Related to Figure 6.. Probabilities for the target and its neighbors were computed by applying the classifiers trained without target peptides and their neighbors. See the legend of Figure 6 for further information.

### Supplementary Tables

**Table S1.** Variance inflation factors (VIFs) of the features used in the multivariate regression analysis against experimentally determined TCR affinities..

| Feature | VIF |
| --- | --- |
| sCPP_BETM990101inv_Skew_4 | 1.25 |
| sCPP_KESO980102inv_Kurt_3 | 1.11 |
| sCPP_KOLA930101_Skew_4 | 1.29 |
| sCPP_SIMK990104_SD_3 | 1.14 |
| sCPP_VENM980101inv_Min_3 | 1.06 |

## Supplementary Datasets

**Supplementary Datasets Data S1. (separate file)**

Human, primate, and rodent T cell epitope dataset.

**Data S2. (separate file)**

TCR-peptide affinity and structure dataset.

**Data S3. (separate file)**

Feature importance estimates.

**Data S4. (separate file)**

Immunogenicity scores.

**Data S5. (separate file)**

A list of single amino acid mutant pairs identified.

**Data S6. (separate file)**

A list of AAIndex amino acid pairwise contact potential scales.

